# Coupled Transcriptomic and ECM-Mechanical Remodeling Reveal Mechanotransductive Pathways in Spinal Cord Injury

**DOI:** 10.64898/2026.07.02.736147

**Authors:** Elif Ertugral, Vasanti Dhakate, Rounak Pokharel, Gowsia Banu Shaik, Jessica Onyak, Peng Jiang, Chandrasekhar R. Kothapalli, Nic D. Leipzig

## Abstract

Spinal cord injury (SCI) leads to the formation of a chronic scar composed of glial and fibrotic components that severely restrict neural regeneration and functional recovery. While the scar composition has been widely studied, the spatiotemporal evolution of tissue mechanics and the role it plays in regulating the post-injury responses remain poorly understood. Here we present an integrated mechanobiological and multi-omics analysis of spinal cord remodeling following a severe thoracic contusion injury. Using nanoindentation and viscoelasticity measurements taken via atomic force microscopy (AFM), we demonstrate that SCI induces a dynamic mechanical response characterized by rapid tissue softening during the acute phase reaching a minimum at one-month post-injury, followed by progressive stiffening associated with chronic scar maturation at six months. Bulk RNA sequencing reveals that early mechanical softening coincides with strong activation of inflammatory and matrix-degrading pathways whereas chronic stiffening correlates with upregulation of collagen synthesis, extracellular matrix (ECM) organization and fibrotic remodeling pathways. Concurrently, mechanotransduction regulators exhibit temporally coordinated activation, indicating that cells dynamically sense and respond to evolving mechanical cues. Viscoelastic analysis further shows that chronic scar tissue exhibited increased stiffness and prolonged relaxation dynamics, reflecting dense collagen deposition and proteoglycan accumulation that reinforces a mechanically restrictive microenvironment. Together, these findings establish that the post-injury scar represents a dynamic mechanobiological system in which the evolving tissue mechanics, viscoelasticity and mechanotransduction collectively regulate ECM remodeling, resulting in regenerative failure. This study provides a comprehensive mechanobiological framework for SCI progression and highlights the opportunities for mechanically informed therapeutic strategies aimed at modulating scar mechanics to promote tissue repair.

**Graphical Abstract:** 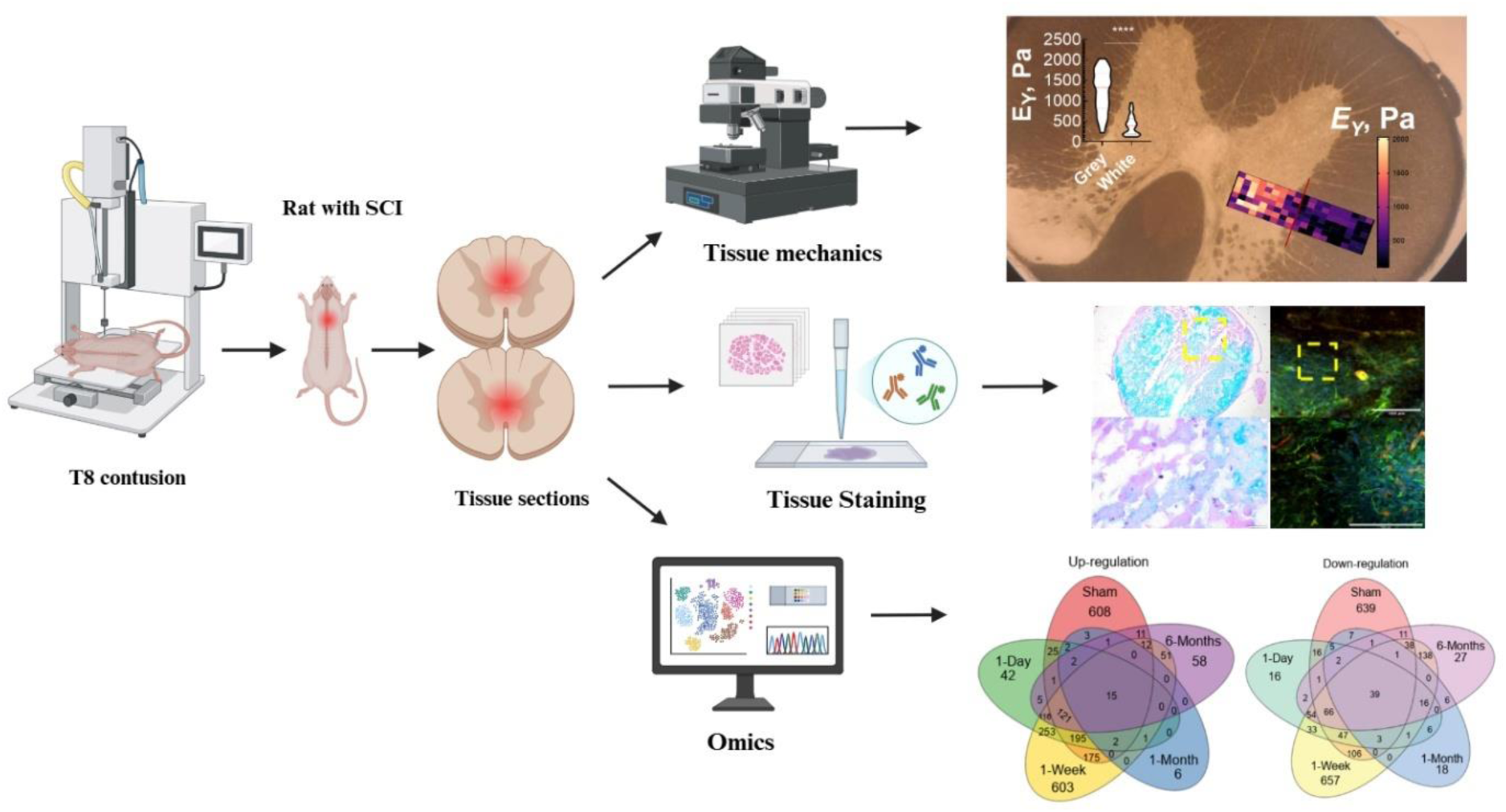

## Introduction

Spinal cord injuries (SCI) affect nearly 300,000 people in the U.S. and over 15 million globally, highlighting the urgent need for better treatments [1]. Despite advances in acute care, clinically proven pharmacological or surgical intervention does not exist that can reliably restore cellular processes and function following SCI [2, 3]. Regenerative strategies remain largely experimental, hampered by the complexity of spinal cord biology and the hostile post-injury microenvironment that actively resists repair, underscoring the need to better understand the post-injury microenvironment to better inform next-generation therapeutic strategies [4].

SCI progression occurs through a biphasic cascade involving primary and secondary injury mechanisms [5], which makes it difficult to treat. The primary injury post-trauma results in immediate mechanical disruption of neural tissue, blood vessels, and cellular structures [6]. This triggers a series of secondary injury processes characterized by inflammation, oxidative stress, excitotoxicity, and extensive remodeling of the extracellular matrix (ECM) [7]. Such secondary events progressively exacerbate tissue damage and culminate in the formation of a complex scar tissue at the lesion site [8]. Regenerative failure following SCI has been attributed to the formation of the glial scar, which arises from reactive astrogliosis and is associated with the upregulation of neuronal inhibitory ECM molecules, particularly chondroitin sulfate proteoglycans (CSPGs) [9]. Although therapeutic strategies targeting CSPGs have demonstrated partial success in promoting axonal growth, they have not been sufficient to restore robust neural regeneration, suggesting that additional structural and molecular barriers contribute to the inhibitory microenvironment in the injured spinal cord [10, 11]. This recommendation underscores the need to assess spatial and temporal dynamics to modulate the inhibitory microenvironment and identify optimal therapeutic windows for promoting axonal regeneration.

Recent studies indicate that the lesion site comprises not only a glial scar, but also a substantial fibrotic scar component [12]. This fibrotic core is formed primarily by infiltrating fibroblast-like cells, perivascular stromal cells, and pericyte-derived cells that deposit structural ECM proteins such as collagen, fibronectin, and laminin [13–15]. Surrounding this core, reactive astrocytes produce proteoglycans and glycosaminoglycans (GAGs), creating a glial-rich ECM environment that further shapes cellular behavior and tissue remodeling, while also inhibiting axonal growth [9]. Together, these persistent fibrotic and glial components generate a heterogeneous scar architecture with distinct biochemical and structural properties.

Despite these established findings, the spatial mechanical phenotype of the scar at both acute and chronic stages remains poorly defined, and the shifting balance between elastic and viscous ECM components during the transition from acute inflammation to chronic scar formation remains unresolved. A comprehensive characterization of this mechanobiological remodeling requires approaches capable of simultaneously resolving tissue-scale mechanical changes, genome-wide transcriptional programs, and protein-level ECM dynamics. While bulk RNA sequencing provides the primary molecular readout of injury-induced gene expression changes, transcript abundance alone does not fully capture the post-injury landscape particularly for ECM-associated proteins whose levels are regulated post-translationally and whose structural insolubility can limit mass spectrometry detection [16]. To provide complementary translational context, integrated proteomics alongside transcriptomics, focusing on pathway-level enrichment to identify shared and divergent molecular signatures across post-injury timepoints are required. Together, these modalities could offer a more complete view of injury progression than either approach alone [17, 18].

In this study, we present an integrated map of how the mechanical landscape of the spinal cord evolves in parallel with molecular remodeling after injury. We elucidate and correlate tissue-scale mechanical properties, structural architecture, and mechanochemical signaling across key post-injury phases using atomic force microscopy (AFM) for spatiotemporal characterization. By identifying sub-cellular responses to altered local biomechanics, we provide a multi-disciplinary and comprehensive analysis of glial scar micromechanics, ECM composition, and associated cell behaviors. Such centering of tissue mechanobiology represents a paradigm shift in CNS repair strategies and may help advance efforts to overcome the limited regenerative capacity of the adult CNS.

## Results and Discussion

### Injury induces spatiotemporal biomechanical changes in spinal cord tissues

To characterize spinal cord tissue mechanics, AFM nanoindentation was performed on naïve and sham rat samples to quantify the linear elasticity of white and grey matter. Contact-mode force–indentation curves (**Supplementary Fig. 1**) were fitted using a modified Hertz model to derive the Young’s modulus (E_Y_). The average E_Y_ of grey matter (1223 ± 497 Pa) was 3-fold higher than that of white matter (396.7 ± 200 Pa) in naïve tissues (*p* < 0.0001; **Supplementary Fig. 2**), whereas the E_Y_ of grey matter (509.7 ± 262.4 Pa) was 1.2-times higher than that of white matter (402.9 ± 165.5 Pa; *p* = 0.1063) in sham tissues (**Supplementary Fig. 3**). Although there is statistically no significance in E_Y_ between grey and white matter in sham tissues (*p* = 0.1063), the sharp drop in elasticity of grey matter compared to naïve tissue (*p* < 0.0001) underscores that a trauma and resultant tissue level changes still occur despite not directly interacting with the spinal cord during laminectomy. Thus, laminectomy appears to compromise the spinal cord’s intrinsic tissue stiffness, thereby distorting the mechanical gradient naturally observed in naïve tissue. Overall, the tissues exhibited average values of approximately 800 Pa for naive and 450 Pa sham, providing a strong foundation for characterizing tissue properties and assessing surgical effects.

Next, the spatiotemporal mechanical heterogeneity of injured rat spinal cord sections was assessed across multiple post-injury time points and three defined regions of interest. Injured tissue sections showed distinct differences in stiffness compared to sham and naïve tissues, with a significant decrease in stiffness within the injury region compared to adjacent and far regions (**Fig. 1C**, *p* < 0.0001, **Supplementary Table 1**). The significant decrease in E_Y_ at the injury region by 1-month (65%, 80%, and 86% drop at 1-day, 1-week and 1-month respectively, **Fig. 1D**; *p* < 0.0001) coincides with the development of a glial-fibrotic scar. A progressive increase in stiffness at 6-months (53% drop; Fig. 1D) corroborates the dynamic nature of scar formation, indicating a transition to a stabilized chronic scar. Because the glial scar is generally considered mechanically stiffer than intact tissue [19, 20], it is widely viewed as a physical barrier that impedes neural regeneration in the CNS [21]. Our findings align with those of Moeendarbary et al. who reported glial scar softening of rat spinal cord after 9-days post-injury with persistence through 3-weeks, while Jin et al., observed initial acute softening 3-days post-injury, transient to the baseline by 6-weeks and followed by increased stiffness during the chronic phase by 12-weeks post-injury [22, 23].

**Figure 1:**
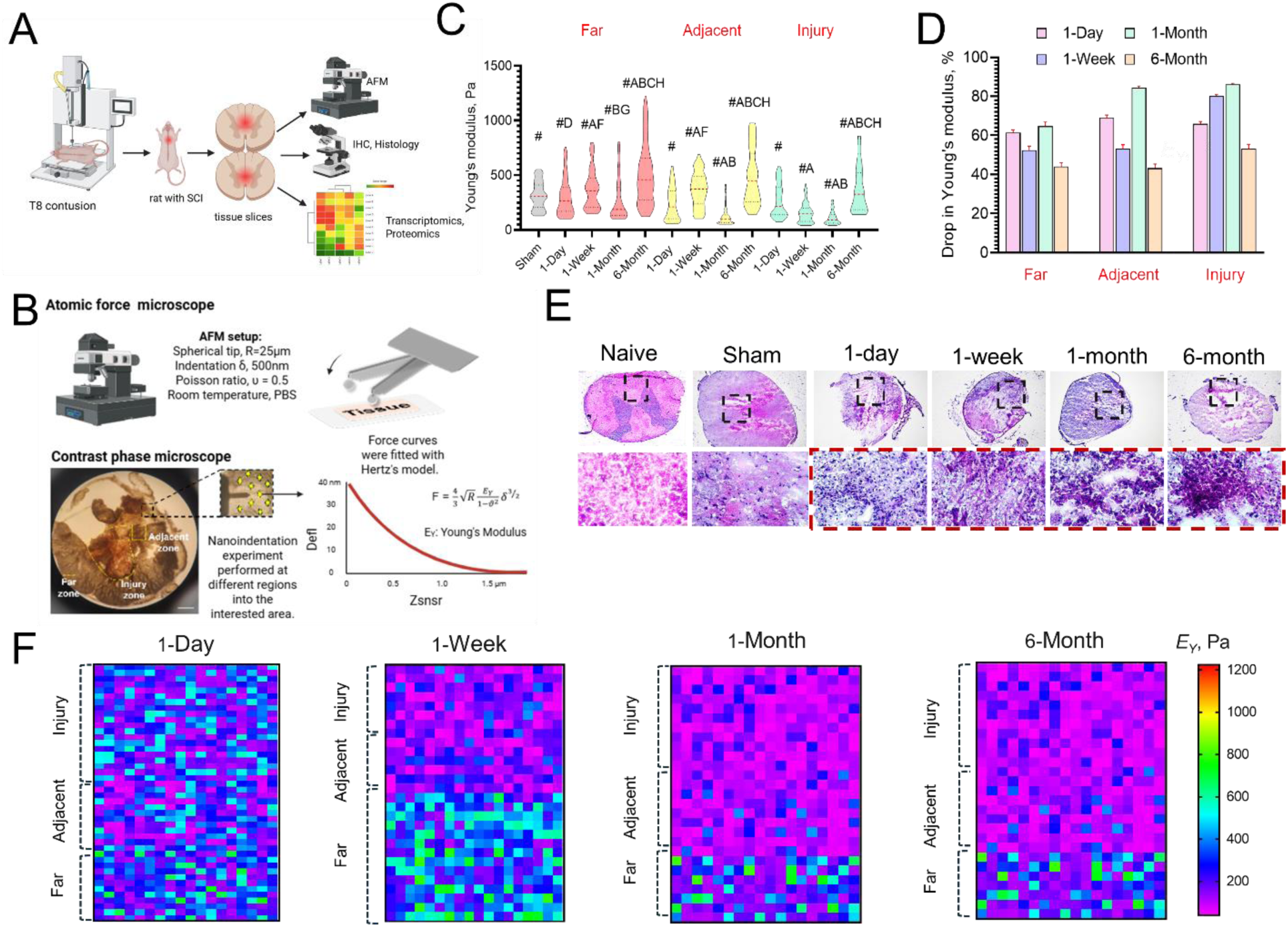
Biomechanical characterization of injured rat spinal cord using AFM. (**A**) Overview of the experimental workflow. (**B**) Schematic of the nanoindentation workflow on a spinal cord hemi-section, highlighting the three regions of interest selected for spatiotemporal analysis of post-injury mechanical changes. (**C**) Spatiotemporal changes in E_Y_ of rat spinal cord tissues – three different regions (far, adjacent, and injury) and four post-injury time points (1-day, 1-week, 1-month, and 6-month) were tested. Data shown as mean ± SEM (108 ≤ n ≤ 360 data points; N=6 animals for all data). Statistical significance was calculated by one-way ANOVA with Tukey’s *post hoc* test to independently evaluate significant spatiotemporal differences; subsequent uppercase letters represent differences between time points, A: vs 1-day, B: vs 1-week, C: 1-month, D: 6-month, within each regions while letters E-H represent significant differences corresponding to the time points between regions. Groups with the same letter are not significantly different (*p* < 0.05). Statistical significance compared to naïve tissue was determined using an unpaired Student’s t-test (# indicates *p* < 0.0001, data in **Supplementary Table 1**). (**D**) Regional relative drop in E_Y_. (**E**) Representative H&E-stained images of the tissues at various timepoints; purple shows cell nuclei, and pink labels the ECM/cytoplasm. The red box indicates the injury region. Scale bars: 200 µm (low magnification) and 50 µm (high magnification). (**F**) Representative heat maps of E_Y_ at 1-day, 1-week, 1-month, and 6-month time points, each shown at three different regions on the injured T8 tissue.

H&E staining (**Fig. 1E**) revealed progressive loss of tissue integrity after injury, shifting from the organized cellular architecture of naïve and sham tissues with distinct demarcation of grey and white matter, to severe structural disruption and the formation of a dense, acellular lesion core by 6-months, consistent with mechanical stiffening. These observations point to a model in which pronounced tissue softening dominates the acute phase and radiates outward from the injury site, followed by substantial stiffening as the lesion matures into a chronic scar. Overall, the distinct mechanical profiles observed across the four post-injury time points (**Fig. 1F**) indicate that elasticity decreases acutely and propagates outward from the injury core with time, while chronic scar formation ultimately stiffens the lesion core.

### Transcriptional and translational evolution post SCI correlates with biomechanical changes

To further examine the molecular signaling cascade that parallels evolving biomechanical cues after SCI, we analyzed temporal transcriptomic changes between the acute and chronic phases of SCI. Differentially expressed genes (DEGs) in sham, acute, and chronic tissues were compared with those in naïve controls, and statistical analysis of normalized read counts was used to quantify differences in DEG expression across groups. The resulting overlaps were visualized using Venn diagrams (**Fig. 2A**). Compared with naïve tissue, we observed a pronounced transcriptional shift at 1-week post-injury, with 603 distinct genes upregulated and 657 downregulated during the subacute phase (1260 distinct total DEGs). Gene expression changes were minimal at 1-month post-injury (six distinct total DEGs), reflecting a sharp decline after the early peak. By 6-months, however, a secondary wave of gene upregulation emerged, coinciding with the progressive mechanical stiffening of the chronic scar. The mechanical stiffening observed at 6-months may reflect a long-term consequence of the intense transcriptional activity initiated at 1-week post-injury. The marked reduction in gene expression at 1-month likely signifies the resolution of the acute response, whereas the renewed surge at 6-months suggests ongoing chronic remodeling or fibrotic processes.

**Figure 2.**
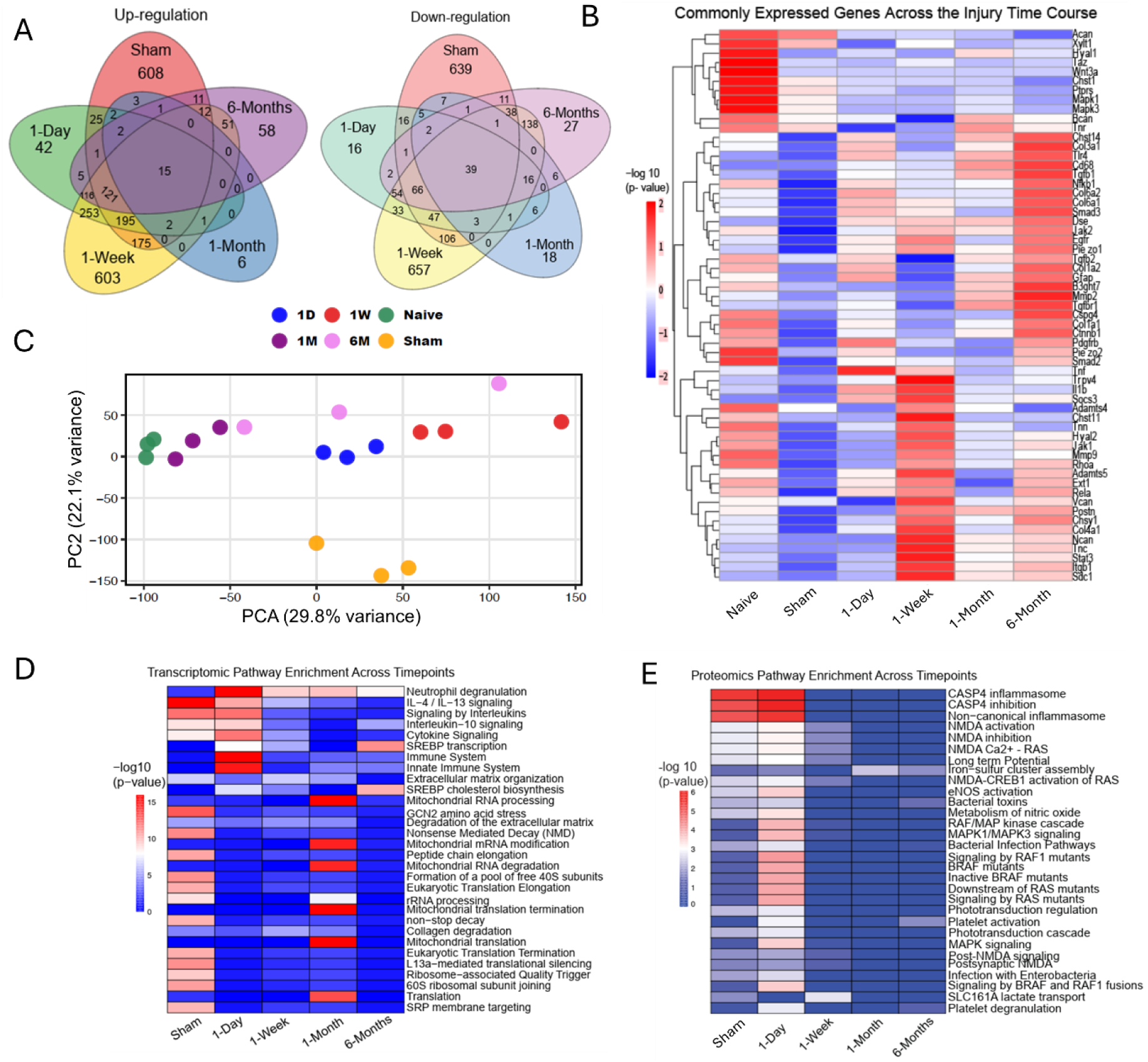
Multi-omics characterization of temporal molecular changes following spinal cord injury. (**A**) Venn diagrams illustrate the overlap of significantly upregulated and downregulated genes across post-injury time points relative to naïve tissue. (**B**) Heatmap showing averaged expression levels of injury-associated genes across naïve, sham, and post-injury samples (log_10_-normalized counts). (**C**) Principal component analysis (PCA) demonstrating temporal separation of transcriptomic profiles after injury; each point represents one biological replicate. (**D**) Significantly enriched pathways identified from differentially expressed genes at each time point (−log_10_ p-value). (**E**) Proteomics-derived enriched pathways highlighting time-dependent molecular remodeling following injury. For all data, n = 6 animals per group.

Consistent with the temporal trends observed with our findings, Shi et al. reported increased transcriptional activity at the acute phase (1-day post injury, 1707 DEGs) followed by a peak at the subacute phase (6-days post-injury, 6590 DEGs) and reduced transcriptional activity at 28 days with 3499 DEGs [24]. Although the absolute number of DEGs differed, both studies demonstrate an early transcriptional surge followed by attenuation at later stages. Similarly, Duan et al. confirmed ongoing chronic remodeling, with elevated immune, defense, and wound responses, for at least 90 days after injury [25]. Compared to these prior studies, our 6-month time point portrays a true chronic phase. The distinct increase in gene expression detected in the sham group may represent compensatory responses to surgical stress in the absence of injury-driven signaling. Accordingly, subsequent analyses focus on comparisons of injured time points relative to naïve tissue, with the full sham dataset shown in **Supplementary Fig. 4**. In the injury groups, the DEGs highlight damage-induced signaling pathways over the sham-related response. Notably, gene upregulation reaches its lowest point at 1 month, with only 6 upregulated and 18 downregulated distinct DEGs when the tissue is mechanically collapsed, indicating a period of transcriptomic silence that may represent a critical window during which subcellular machinery adapts to the loss of structural support. This finding strongly mirrors the recent rodent studies, confirming the temporary suppression of active transcription in cells to adapt to altered ECM biomechanics [26].

We next examined genes commonly expressed across post-injury time points to correlate molecular changes with the mechanical shifts observed in AFM measurements (**Fig. 2B**). Inflammatory and degradative genes (*IL1β, STAT3, MMP9, ADAMTS4*) were strongly upregulated at 1-week, corresponding to the marked reduction in E_Y_ during the subacute phase [27–29].Our findings are consistent with those of Mun et al. who used a rat contusion model and reported a peak inflammatory response at 1-day post-injury, characterized by dominant interleukin signaling and substantial collagen degradation by 3-months [30]. In contrast, robust upregulation of fibrotic genes (*COL3A1, COL6A1, COL6A2, COL1A1, COL1A2*) emerged in the chronic phase, aligning with increased stiffness and extensive ECM remodeling [12, 31]. Gong et al. corroborated these trends using RNA-seq at 1, 4, and 7-days after SCI, identifying strong enrichment of TNF and cytokine–cytokine receptor interaction pathways that drive the early shift toward active ECM degradation during the first week [32]. Altered expression of key mechanotransduction genes (*PIEZO1/2, TAZ*) indicates active cellular sensing and response to changes in the biomechanical environment [33, 34]. Principal component analysis reveals a clear trajectory among PC1 compared to naïve, highlighting high genomic expression at the 1-week time point. Interestingly, the sham group is clearly distinct from all injury conditions among PC2, indicating a distinguished biological profile reflecting the surgical procedure itself (**Fig. 2C)**. Overall, the acute phase is characterized by a simultaneous surge in *TGFβ1* (a driver of chronic scar formation) and *MMP* family genes (mediators of matrix degradation), which together underlie the rapid mechanical softening observed at 1 week. In contrast, the chronic upregulation of collagen genes (*COL1A1, COL3A1*) and inhibitory CSPGs (*ACAN, VCAN, NCAN*) provide genetic evidence for the stiff, viscous barrier that defines the 6-month scar.

Pathway enrichment analysis of the gene expression data (**Fig. 2D**) showed that injury rapidly induced expected strong immune and cytokine signaling responses at 1-day, followed by pronounced MMP-mediated matrix degradation at 1-week. These early events then transition into sustained ECM organization and collagen synthesis, which dominates the chronic phase and defines the long-term scar phenotype at 6-months. Proteomic pathway enrichment analysis (**Fig. 2E**) revealed that protein-level inflammatory signatures were most prominent at 1-day post-injury, with strong enrichment of CASP4-mediated inflammasome activation, NMDA receptor signaling, and MAPK/ERK cascades at the acute timepoint. This proteomics signal progressively declined across subsequent time points, consistent with the resolution of the acute inflammatory response and a transition toward more structurally driven remodeling programs. The dominance of inflammatory and neuronal signaling pathways in the proteomic data, rather than ECM organization pathways likely reflects the known underrepresentation of insoluble fibrillar ECM proteins, such as collagens in standard mass spectrometry workflows, despite their robust transcriptional upregulation [16]. Together, the transcriptomic and proteomic profiles converge on the same biological trajectory: a peak inflammatory surge at 1-day that sees the progressive ECM remodeling, with each modality capturing complementary aspects of this transition at the gene-expression and protein-activity levels, respectively.

### Spatiotemporal evolution of viscoelasticity in spinal cord tissues

The ECM is a major structural component of tissue, composed of macromolecules arranged in a complex 3D network. Under stress, this network undergoes protein unfolding, polymer disentanglement, and rupture of weak bonds, producing the characteristic time-dependent viscoelastic behavior of tissues [35]. Crosslinked collagen fibers dissipate energy by breaking noncovalent interactions [36], while other ECM macromolecules dissipate stress through chain rearrangement and matrix flow. Interstitial fluid movement also contributes to viscoelasticity. Cells actively sense and respond to matrix deformation, while the ECM gradually relaxes internal stress under constant strain – a process known as stress relaxation. This molecular rearrangement helps maintain tissue equilibrium and protects against damage. Matching the stress-relaxation timescales of biomaterials to those of native tissues is crucial for regenerative medicine applications [37, 38].

To assess how the spinal cord tissue responds to mechanical loading and to quantify its viscoelastic properties to better understand ECM remodeling, we performed stress-relaxation tests on naïve, sham, and injured fresh samples (**Figure 3**). For each condition, at least 20 force curves were collected using step indentation, during which the AFM probe indented the tissue and held a constant depth for a 5-second dwell period to capture force decay. Longer dwell times yielded no additional benefit to measurements. Data was collected from both white and grey matter in naïve and sham tissues, and from injury-core, adjacent, and far-field regions in injured cords to resolve spatiotemporal changes after SCI. We quantified three key viscoelastic parameters: the instantaneous modulus (E_0_), reflecting the purely elastic stiffness at the moment stress is applied; the infinite modulus (E_inf_), representing the elastic response after relaxation is complete; and the relaxation time (*τ*_α_), indicating the rate at which the material transitions from its initial to equilibrium state. Values of *τ*_α_ were extracted from the force-decay profile, while those of E_0_ and E_inf_ were derived from the initial peak and final force plateau, respectively. Frequency-dependent properties – storage (G′) and loss (G″) moduli – were obtained by fitting the data to a Standard Linear Solid (SLS) model (**Fig. 3A**).

**Figure 3:**
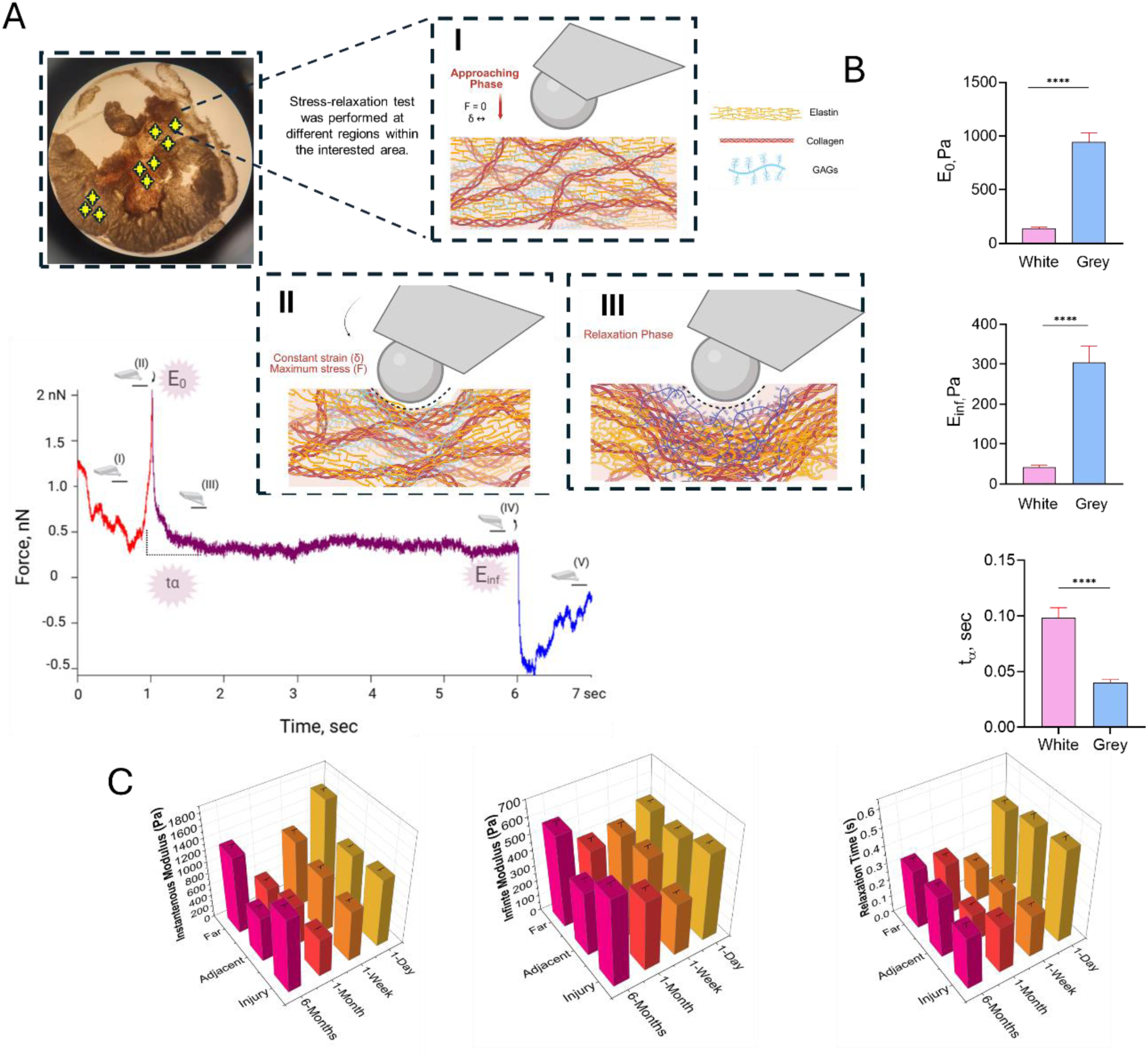
Viscoelastic characterization of injured rat spinal cord. (**A**) Stress-relaxation testing used to quantify spinal cord tissue viscoelasticity: (I) Initial pause with the probe positioned above the sample, followed by approach. (II) Constant-indentation phase at the maximum applied force. (III) Relaxation phase, during which force decays under constant strain. (**B**) Viscoelastic properties of naïve adult rat T8 spinal cord tissue, including instantaneous modulus (E_0_), infinite modulus (E_inf_), and relaxation time (t_α_). Data was presented as mean ± SEM, calculated from the force-time plots obtained using AFM, with model fit in Viscoindent. Statistical significance was determined using an unpaired Student’s t-test (79 ≤ n ≤ 94 points for E_0_, 65 ≤ n ≤ 88 points for E_inf_, 63 ≤ n ≤ 82 points for t_α_, **** indicates *p* < 0.0001). (**C**) Viscoelastic characteristics (E_0_, E_inf_, t_α_) of injured adult T8 tissues, at acute and chronic time points. Statistical significance was determined using one-way ANOVA (22 ≤ n ≤ 178 points for E_0_, 28 ≤ n ≤ 169 points for E_inf_, 35 ≤ n ≤ 181 points for t_α_; **** indicates *p* < 0.0001; data in **Supplementary Table 2**) with *post hoc* Tukey’s HSD tests. N=3 animals for all data.

Substantial mechanical heterogeneity was noted in naïve tissue, with grey matter exhibiting significantly higher E_0_ (945.8 ± 84 Pa) and E_inf_ (304.7± 40.8 Pa) compared to white matter (142 ± 9.8 Pa and 42.4 ± 5 Pa, respectively; *p* < 0.0001; **Fig. 3B**). Conversely, white matter exhibited a longer *τ*_α_ (0.099 ± 0.008 s) and greater viscous response than the more elastic dominated grey matter (0.04 ± 0.003 s; *p* < 0.0001). As in naïve tissue, sham tissues maintained biomechanical heterogeneity, with grey matter exhibiting higher E_0_ and E_inf_ than white matter (**Supplementary Fig. 5**). Conversely, grey matter showed a slightly prolonged *τ*_α_ (0.098 ± 0.016 s) than the white matter (0.076 ± 0.013 s), showing the biomechanical profile was heavily influenced by inflammation response and was associated with edema rather than the intrinsic properties (*p* < 0.0001 for E_0_ and E_inf_; *p* = 0.3436 for *τ*_α_).

Using the same protocol, we next performed stress-relaxation testing on injured fresh spinal cord sections (**Fig. 3C**). Analysis showed that injury induces pronounced mechanical softening, particularly in E_0_, which peaked at the injury site at 1-day (1181.6 ± 24.5 Pa), declined sharply by 1-week (889.2 ± 42.2 Pa), and continued to fall to reach the lowest point at 1-month (663.7 ± 29.8 Pa). By 6-months, mechanical recovery was evident, with stiffness rising to 1280.7 ± 68.5 Pa (*p* < 0.0001 for 1-week and 1-month vs 1-day, *p* < 0.01 for 6-month vs 1-day, *p* > 0.1 (ns) for all other cases).

Our findings align with those of Jin et al. who observed a biphasic mechanical response in the injured rat thoracic spinal cord – an initial softening that reached its minimum at 1-month post-injury, followed by pronounced stiffening by 6-months. They also agree with trends reported by Cooper et al., attributing this chronic stiffening to fibrosis and ECM deposition [31, 39]. Consistent with these studies, E_inf_ in our data follows a similar trajectory (1-day: 503.8 ± 17.4 Pa; 1-week: 425.6 ± 31.4 Pa; 1-month: 412.6 ± 25.9 Pa; 6-month: 581.4 ± 33 Pa), producing a sustained reduction in energy dissipation (E_0_–E_inf_) through 1-month post-injury (*p* < 0.0001 1-week vs 1-day, *p* < 0.01 for 1-month vs 1-day, *p* = 0.0382 for 6-month vs 1-day, *p* < 0.01 for 1-month vs 1-week, *p* = 0.0174for 6-month vs 1-week and *p* = 0.9767 (ns) for 6-month vs 1-month). Notably, this gap increases sharply at 6-months, where E_inf_ becomes disproportionately large relative to E_0_, suggesting a dominant contribution from a dense, crosslinked collagen network [40–42].

This disorganized matrix reduces hydraulic permeability and energy dissipation, resulting in a longer *τ*_α[43]_. In the acute phase, *τ*_α_ is markedly prolonged (1-day: 0.53 ± 0.02 s), reflecting inflammation, edema, and ECM disruption (*p* = 0.0219 vs 1-week and *p* = 0.0123 vs 1-month; *p* = 0.3660 (ns) vs 6-month). It decreases substantially in the subacute (1-week: 0.24687 ± 0.01 s) and intermediate (1-month: 0.27 ± 0.01 s) phases, consistent with a progressive loss of GAGs and interstitial fluid. This downward trend reverses in the chronic phase (6-month: *τ*_α_ = 0.3 ± 0.02 s; *p* < 0.01 vs 1-week and 1-month), where *τ*_α_ increases again, indicating a transition from acute ECM degradation to functional remodeling, including GAG replenishment and fluid entrapment within the disorganized collagen network [22, 31, 44]. Elevated *τ*_α_ was found to be driven by increased tissue hydration profile, suggesting that mature scar has a high capacity for water content within its dense proteoglycan matrix (**Supplementary Fig. 6**). Consequently, injured tissue retains mechanical loads for longer durations, further restricting interstitial fluid flow relative to healthy tissue.

Beyond the injury epicenter, spatiotemporal analysis of the adjacent and far regions revealed a mechanical disturbance that propagated outward from the lesion but diminished with increasing distance from the injury site. In the adjacent region, the E_0_ followed a trajectory similar to the injury site, showing progressive softening from 1-day (804.7 ± 35.2 Pa; *p* < 0.0001 vs 1-week and 1-month; *p* > 0.05 vs 6-month) through 1-month (281.2 ± 52.1 Pa; *p* > 0.05 vs 1-week; *p* < 0.0001 vs 6-month), followed by partial stiffness recovery at 6-months (843± 142.5 Pa). The E_inf_ in the adjacent region exhibited corresponding trends: the gap between E_0_ and E_inf_ narrowed through 1-month (E_inf_: 60.3 ± 17.3 Pa) and widened again by 6-months (E_inf_: 352.1± 66.8 Pa), reflecting evolving energy dissipation capacity as fibrotic remodeling progresses into the perilesional tissue (*p* < 0.0001 for 1-week and 1-month vs 1-day; *p* < 0.0001 for 1-week and 1-month vs 6-month; *p* > 0.05 for all other cases).

The region farthest from the lesion epicenter demonstrated the most modest mechanical perturbations across all timepoints, yet measurable deviations from naïve tissue values were still evident (*p* < 0.0001 vs 1-day; *p* < 0.01 vs 6-month; *p* > 0.05 for all other cases) – particularly, at 1-week (E_0_: 886.2 ± 134.1 Pa) and 1-month (E_0_: 602.3 ± 139.1 Pa) – suggesting that secondary injury-mediated ECM disruption and inflammatory signaling extend beyond the lesion core. In the adjacent region, *τ*_α_ mirrored trends at the injury site, remaining prolonged at 1-day (0.51 ± 0.03 s), due to edema and acute ECM disruption, then shortening through the subacute and intermediate phases as ECM and interstitial fluid content declined, followed by a modest rebound at 6-months (0.35 ± 0.02 s), consistent with the onset of chronic remodeling (*p* = 0.0138 for 1-day vs 1-week; *p* > 0.1 for all other cases). In contrast, the far region displayed comparatively stable *τ*_α_ across all timepoints (*p* < 0.0001 for 1-day and 1-month vs 6-month; *p* > 0.05 for all other cases), approaching values characteristic of naïve tissue (white matter: 0.09 ± 0.008 s, grey matter: 0.03 ± 0.03 s), indicating that viscoelastic disruption attenuates with increasing distance from the lesion.

These spatially graded mechanical changes across injury, adjacent, and far regions collectively demonstrate that the mechanical consequences of SCI extend well beyond the lesion core, representing a distributed mechanobiological disturbance in which the injury epicenter represents the most severe manifestation of a broader tissue-wide remodeling cascade. Consistent with this interpretation, Saxena et.al. measured *τ*_α_ and viscosity in the injured rat spinal cord at 2- and 8- weeks post-injury and reported that injured tissue exhibits a longer *τ*_α_ and higher viscosity, even in the presence of a reduced E_Y_ [45]. Moreover, scar tissue maintains mechanical load on growing axons rather than yielding, thereby influencing neural stem cell fate: prolonged *τ*_α_ promotes neural differentiation, whereas shorter *τ*_α_ drives astrocytic differentiation through RhoA signaling [46]. Similarly, substrates with shorter *τ*_α_ enhance neural maturation and gene expression but impair oligodendrocyte differentiation, potentially suppressing regeneration via mechanotransduction pathways [47, 48]. Collectively, these findings underscore that the scar is not merely a chemical impediment but a mechanical barrier whose altered viscoelastic dynamics instruct resident cells to adopt non-regenerative phenotypes.

Despite extensive literature on CNS mechanobiology, our understanding of mechanical alterations caused by injury remain relatively sparse [49]. To date, very few studies have considered the combined glial and fibrotic scar’s architecture and its structure-function properties as a potential contributing factor to SCI pathology [50, 51]. While studies have shown that substrate mechanical properties direct CNS cells *in vitro* [52, 53], the altered mechanical properties of the fibrotic-glial scar microenvironment and causation are mostly ignored as a factor of SCI pathology. Current therapeutics that target inhibitory CSPGs or immune response do not result in complete recovery, indicating that additional factors of the glial scar, including other chemical and structural properties, need to be considered [51]. The mechanical and biochemical cascade after primary injury still requires spatiotemporal evaluation to understand the recovery mechanism [49].

### Injury-induced biomechanical alterations regulate cell fate via mechanotransduction

ECM composition differentially shapes the mechanical properties of the injured spinal cord. Proteoglycans, GAGs, and hyaluronic acid (HA) impart highly hydrated and viscous characteristics to neural tissue, whereas fibrillar collagens and other structural proteins provide elastic resistance [54]. A key but often overlooked consequence of this evolving ECM heterogeneity is its influence on the local mechanical environment. ECM mechanics critically regulate cellular behavior through mechanotransduction pathways involving integrin-mediated adhesion, cytoskeletal remodeling, and downstream cascades such as focal adhesion kinase and YAP/TAZ, collectively modulating inflammation, ECM synthesis, and tissue remodeling throughout SCI progression [49, 55, 56].

In the acute phase (first 72-h), inflammation and edema occur as protective responses that activate resident cells (e.g., microglia, astrocytes), leading to increased ECM deposition and altered tissue mechanics [57]. Cells sense these mechanical changes through integrin-binding sites, initiating signaling cascades in which mechanotransduction pathways ultimately inhibit regeneration [58, 59] (**Fig. 4A**). Although this response begins as a protective mechanism, it evolves into a maladaptive process that self-propagates to restrict repair. To identify indicators of mechanotransduction pathway activation, we examined cellular responses to altered tissue mechanics and associated protein secretion. Mechanotransduction-related gene expression was visualized in a heatmap, revealing temporal patterns of up- and downregulation across key pathway components (**Fig. 4B**). Transcriptomic data indicated that mechanical load signaling peaked during the subacute phase (1-week), marked by elevated expression of the mechanosensory genes Piezo1 and Itgb1. Zhang et al. reported marked upregulation of Piezo1 at the lesion site in a brain compression model, validating our findings. In alignment with this, Baker and Hagg showed injury-induced spatiotemporal dynamics of integrin β1 heterodimers, expression patterns centered on the 3–7-days window around the lesion site in rats [60]. This surge in mechanosensitive signaling rapidly drives cellular responses to ECM disruption, extending through 1-month, after which expression declines, reflecting downregulation of mechanotransduction activity.

**Figure 4:**
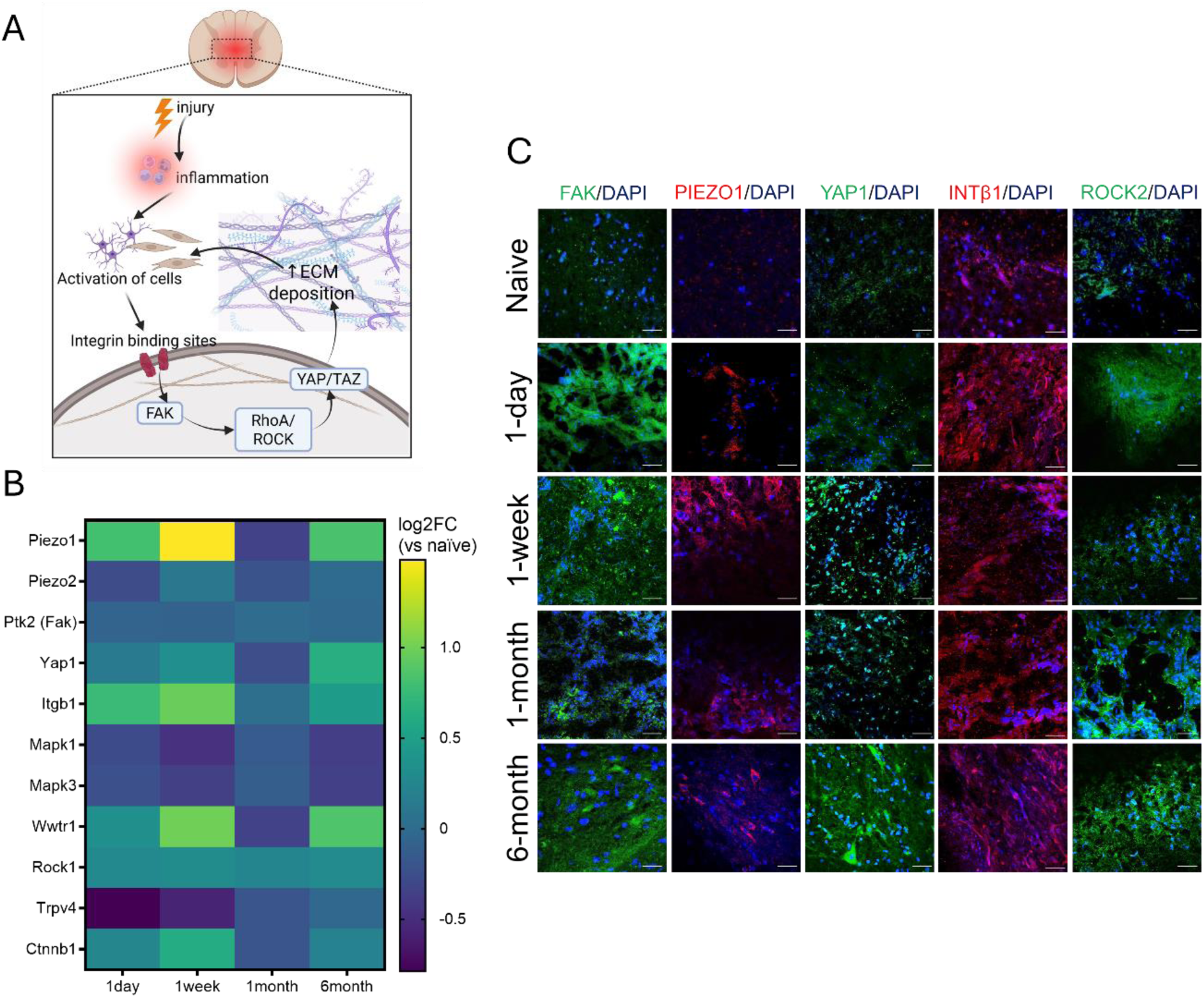
Post-injury ECM remodeling activates mechanotransduction pathways. (**A**) Schematic illustrating injury-induced ECM remodeling and subsequent activation of mechanotransduction signaling cascades. (**B**) Expression of genes associated with mechanotransduction pathways in adult rats following SCI, quantified by bulk RNA-seq. FPKM: fragments per kilobase of transcript per million mapped reads. Statistical significance was assessed using one-way ANOVA (N=6 animals per group, ****indicates *p* < 0.0001) with *post hoc* Tukey’s HSD tests. (**C**) Temporal expression of key mechanotransduction-related proteins in naïve tissue and the injured lesion core. Images were acquired at 60× magnification; corresponding low-magnification (4×) images are provided in **Supplementary Fig. 7.** Scale bars: 50 µm.

By 6-months post-injury, the tissue has undergone substantial remodeling driven by chronic scar-associated programs and become mechanically stiffer, as evidenced by the significant and sustained upregulation of Yap1 and Wwtr1 (Taz) in the lesion [61]. In parallel, Rock1 expression remains elevated throughout the 6-month period, consistent with findings from Conrad et al., who reported persistently high levels of RhoA, a direct upstream activator of Rock, particularly in microglia and scar-forming regions for at least 4 months [62]. Additionally, Trpv4 shows a trend toward upregulation by 6 months, supporting the observed increase in relaxation time and indicating enhanced fluid-entrapment capacity (Fig 3C), which enables cells to sense elevated hydraulic pressure within the remodeled matrix [63].

The persistent sub-baseline suppression of Mapk1 and Mapk3 over the 6-month period, despite a transient rebound at 1-month, suggests that mechanical cues increasingly recruit Yap/Taz rather than the classical MAPK/ERK pathway. This temporal shift reflects selective pathway engagement – MAPK/ERK mediates acute mechanosensitive responses, whereas YAP/TAZ becomes the dominant regulator as the tissue undergoes chronic remodeling and stiffening [64].

Overall, the transcriptomic data reveals a distinct biphasic signature that aligns with the mechanical softening observed at 1-month, when tissue stiffness (E_Y_) reaches its nadir, and supports the subsequent recovery of stiffness accompanied by a longer relaxation time (τ_α_) in the chronic scar. Together, these features provide sustained mechanical cues that reinforce pro-fibrotic cell fate by the 6-month mark. These transcriptomic trends were further corroborated by immunofluorescence staining of mechanotransduction markers in naïve and injured spinal cord tissue (**Fig. 4C**). Early-onset expression of mechanical sensors (PIEZO1 and ITGB1), along with increased focal adhesion kinase (FAK), showed elevation within the acute phase (1-day) and remained persistent through the chronic phase (6-month) as a result of SCI. The downstream signaling effector (ROCK2) was upregulated with delayed kinetics, peaking at 1-month, thereby driving the transcriptional cofactor YAP1, which triggers fibrotic tissue remodeling in response to altered microenvironmental biomechanics. Overall, high expression of all probed mechanotransduction markers was observed in the lesion core, further confirming injury-induced activation of mechanosensory pathways (low magnification images in **Supplementary Fig. 7**), whereas sham tissues showed low-intensity staining of the same markers consistent with a homeostatic baseline in the absence of mechanical stress (**Supplementary Fig. 8**). Membrane-localized clustering at 1-day and 1-week post-injury of PIEZO1 and ITGB1 indicates heightened mechanosensitivity during the early phase. β1-integrin was upregulated and spatially engaged in both glial and fibrotic scar-forming cell populations during subacute and chronic SCI (**Supplementary Fig. 9**), supporting its broader role in coordinating multicellular responses to altered tissue mechanics.

Together, the sustained upregulation of PIEZO1, integrin-β1, ROCK1, and nuclear YAP1 localization suggests persistent activation of mechanosensitive signaling throughout SCI progression. These pathways likely contribute to chronic scar maturation by coordinating reactive gliosis, ECM remodeling, and stromal cell activation in response to the evolving mechanical microenvironment. The persistence of these signaling pathways within the chronic lesion supports the existence of a mechanobiological feedback loop in which injury-induced ECM remodeling continuously reinforces non-regenerative cellular states. Collectively, these observations support the existence of a mechanobiological feedback loop (**Fig. 4A**) in which injury-induced ECM remodeling progressively alters tissue mechanics, which in turn reinforces the molecular and cellular programs that sustain chronic scar maturation following SCI. Despite the reduced E_Y_ observed at 1-month post-injury, elevated Rock2 expression together with sustained YAP1 localization suggests that cells continue to generate substantial intracellular tension in response to the mechanically disrupted extracellular environment. By 6 months post-injury, increased Rock2 expression coincided with greater cytoplasmic sequestration of YAP1, potentially reflecting a transition from active remodeling toward stabilization of the chronic scar matrix. This temporal shift further supports the concept that mechanotransduction pathways dynamically evolve throughout SCI progression, initially coordinating acute cellular responses to injury before ultimately contributing to structural consolidation of the chronic scar and restriction of regenerative plasticity.

### Biphasic ECM remodeling drives fibrotic cell fate through mechanotransduction

A scar-forming fate is exacerbated by a temporal shift in mechanotransduction pathways, culminating in a structural overhaul of the spinal cord tissue. The sustained fibrotic phase is marked by extensive ECM remodeling driven by the activation of fibrosis- and scar- associated signaling pathways [65]. To characterize the translational landscape of these pathways, we examined genes across five categories – fibroblast functional activation, TGFβ signaling, mechanosensing, mechanotransduction, and scar maturation – relative to naïve controls (**Fig. 5A**). Fibroblast activation markers (*FAP, PDGFR-Α, ACTA2*) and *TGFβ* signaling genes were broadly upregulated at subacute and chronic timepoints, consistent with sustained fibrotic scar maturation. Src and Axl gene clusters exhibited coordinated upregulation peaking at 1-week, paralleling the mechanosensing and mechanotransduction signatures shown in Fig. 4B. This mechanosensitive interpretation of the injury environment ultimately dictates the long-term fibrotic response driven by TGFβ, culminating in progressive scar maturation [66].

**Figure 5:**
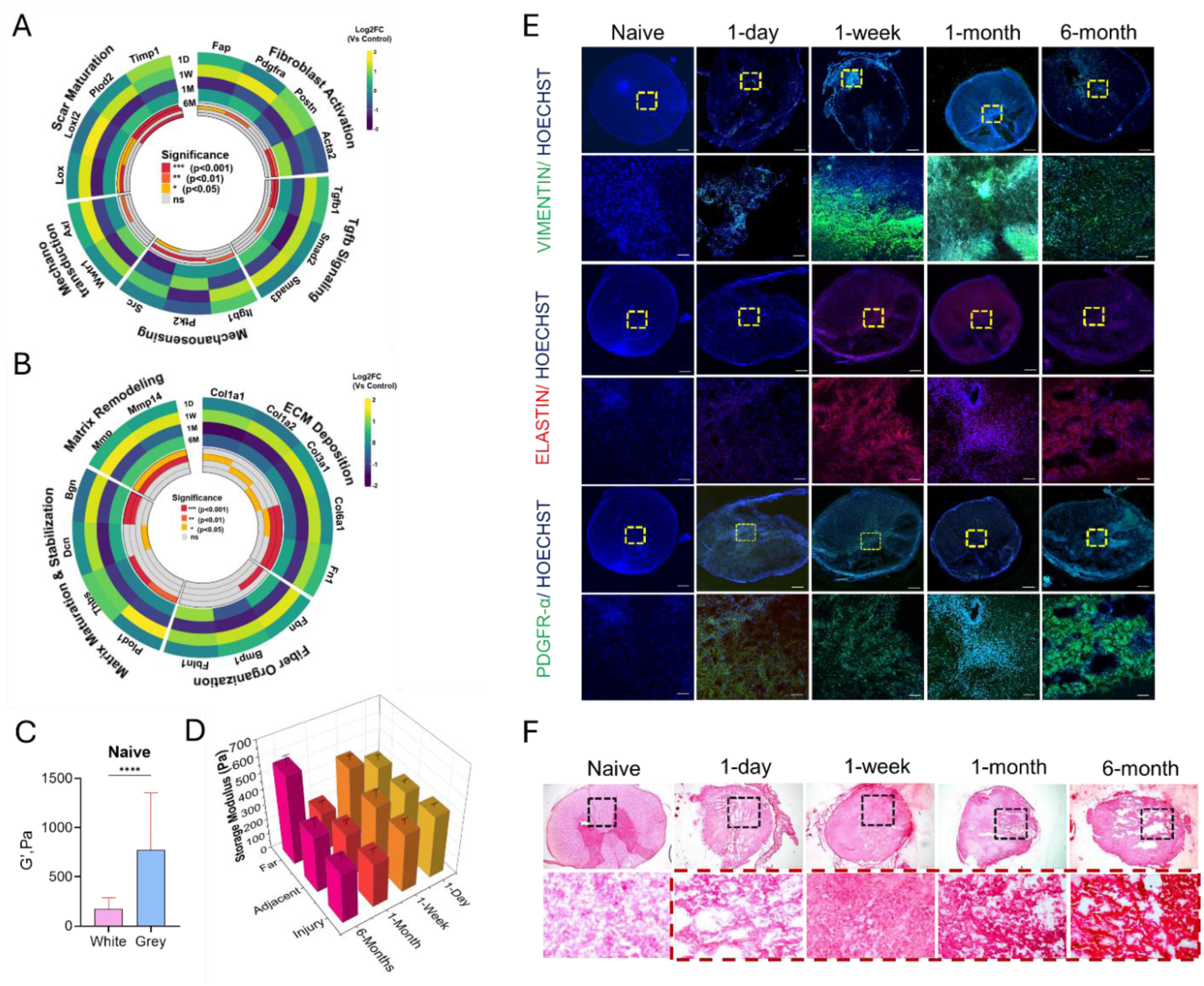
Elastic remodeling following spinal cord injury. Heatmap showing expression of major ECM and matrix remodeling genes (**A**), and fibrosis-associated genes (**B**), across post-injury timepoints. Storage modulus (G′) of naïve tissue (**C**) and injured (**D**) T8 adult rat spinal cord tissue, calculated from AFM-derived force–time curves using Viscoindent model fitting. Statistical significance was assessed using one-way ANOVA with post-hoc Tukey’s HSD tests and unpaired Student’s t-tests (27 ≤ n ≤ 178 data points; N=3 animals for all data; **** indicates *p* < 0.0001). (**E**) Temporal expression of vimentin, PDGFR-α, and elastin proteins, with Hoechst marking nuclei, demonstrating matrix-remodeling activity in naïve tissue and lesion cores. Images were acquired at both high (60×) and low (4×) magnification (**F**) Picrosirius red staining revealed collagen deposition and structural remodeling at the lesion site over time. Scale bars: 200 µm (low magnification) and 50 µm (high magnification).

To assess transcriptional trends in matrix remodeling pathways following SCI, we examined genes across five categories, ECM fiber organization, deposition, remodeling, maturation, and stabilization – relative to naïve controls (**Fig. 5B**). ECM-deposition genes (*COL1A1, COL1A2, COL6A1, FN1*) and matrix-remodeling genes (*MMP2, MMP14*) were broadly upregulated, showing pronounced changes by the 1-month timepoint, indicative of rapid breakdown and rebuilding of structural proteins. Genes associated with matrix maturation and stabilization (*POD1, THBS2, DCN, BGN*) as well as fiber organization (*FBN1, BMP1, FBLN1*) were elevated during the chronic phase, marking the establishment of a permanent fibrotic scar [67]. This translational signature of fibrotic scar formation and ECM remodeling pathways aligns with the biphasic response of the elastic component over the 6-month period. Viscoelastic characterization reveals that beyond the initial softening observed over 1-month (G′: 1-day 409.3 ± 19.6 Pa, 1-week 386.6 ± 34.4, 1-month 293 ± 29.8 Pa), the storage modulus exhibited slightly elevated values at 6-month (333.8 ± 53.7; *p* > 0.05 for all cases), reflecting maturation and stabilization of the injury site. Tissue elasticity shifted over time at the injury site compared to naïve tissue (**Fig. 5C**; white matter: 177 ± 12.1 Pa, grey matter: 771.8 ± 59.6 Pa; *p* = 0.0473 for 1-month, *p* > 0.05 for other cases), suggesting initial softening driven by fiber disorganization and MMP-mediated degradation, followed by a transition to mechanical stiffening, marked by recovery of a higher storage modulus (G′) at the chronic phase. (**Fig. 5D**). Furthermore, the dynamic viscoelasticity of sham tissues exhibited an elasticity-dominated profile, confirming the mechanical consequences of SCI rather than non-specific surgical effect (**Supplementary Fig. 10**). In support of these results, Manesco et al. reported increasingly dense and curved collagen I fiber organization after SCI in mice [12], and Jin et al. also observed increased elasticity along with stiffness following SCI [22].

As with the injury site, spatiotemporal analysis of adjacent and far regions revealed elevated elastic response following SCI, indicating initial softening in the acute phase, followed by mechanical rebound in the chronic phase. In the adjacent site, over the 1-month, diminishing storage modulus (G′) was evident with a trajectory similar to the injury site (1-day: 396.0 ± 21.4 Pa, 1-week: 383.5 ± 54.0 Pa, 1-month: 293.04 ± 33.4 Pa), followed by an elevated G′ at 6-month (370.5 ± 55.9 Pa; *p* > 0.05 vs. all cases). Mechanical perturbations persist in the far regions at all time points, with measurable deviations from naïve tissue values. Interestingly, G″ reached a maximum at 1-month (256.0 ± 24.6 Pa) post-injury, whereas it was closest to that of naïve tissue at 6-month (578.0 ± 66.4 Pa; *p* < 0.001 vs 1-month, *p* = 0.7654 vs 6-month). Together, distortion in elasticity is not confined to the epicenter of injury but propagates outward to the tissue periphery, where mechanical stabilization is incomplete [68].

We further validated the transcriptomic trends through spatiotemporal protein profiling, which revealed elevated expression of vimentin, elastin, and *PDGFR-α* at 1-month, coinciding with the peak of matrix remodeling and the onset of tissue stiffening. Sham tissues exhibited basal staining for these markers (**Supplementary Fig. 11**) confirming that the upregulation of fibrotic markers in test conditions was indeed driven by injury-induced ECM remodeling (**Fig. 5E**). Spatially, vimentin and *PDGFR-α* co-immunoreactivity was concentrated at the lesion core from 1-week post-injury and intensified progressively through the 1-month and 6-month timepoints, consistent with the progressive recruitment and activation of a PDGFR-α+ stromal fibroblasts – the primary cellular driver of fibrotic ECM deposition after SCI [15, 69]. PDGFR-α+ cells initiate the fibrotic program that deposits the dense collagen-rich ECM of the chronic scar. Vimentin co-expression in these PDGFR-α+ cells confirms their mesenchymal, fibroblast-like identity, consistent with evidence that pericyte-derived and meningeal fibroblast populations upregulate vimentin as part of their fibrotic activation program [14].

Elastin staining, most pronounced at the 6-month lesion core, reflects terminal maturation of the fibrotic ECM deposition at chronic timepoint, aligning mechanistically with the elevated E_Y_ (**Fig. 1C**) and prolonged stress-relaxation time captured by AFM (**Fig. 3C**). Critically, the spatial gradient of these three markers – highest at the lesion core and diminishing progressively toward the adjacent and far regions – mirrors the mechanical gradient quantified by AFM nanoindentation, providing direct protein-level evidence that Vimentin+/ PDGFR-α+ stromal cell activation could be the cellular basis of the spatially graded mechanical stiffening observed after SCI. This spatial correspondence between fibrotic cell activation and regional tissue mechanics constitutes a key mechanistic bridge between the molecular and biomechanical dimensions of this study, each independently confirming the same underlying biology.

Picrosirius Red staining confirmed the temporal progression from matrix disorganization to maturation and ultimately stabilization of the fibrotic scar by the 6-month timepoint (**Fig. 5F**). Crucially, the basal reference underscores the gradual collagen deposition and densification observed at the injury site is exclusive to remodeling of SCI and provides a supportive basis to explain the recovery of both G′ and E_Y_ (**Supplementary Fig. 12**).

To further delineate these structural changes, we examined the regulatory mechanisms governed by the non-collagenous matrix, assessing genes involved in CSPG biosynthesis, GAG remodeling, and astrocyte reactivity relative to naïve controls. These analyses highlight glial and proteoglycan-associated pathways that shape ECM dynamics following SCI (**Fig. 6A**). CSPG core proteins (Ncan, Bcan, Vcan) [70–72] and chondroitin sulfate biosynthesis genes (Chsy1, Chst1, Chst11, Chst14) [73]were broadly upregulated at subacute and chronic timepoints, indicating active proteoglycan deposition and enhanced elaboration of inhibitory GAG chains during scar maturation [74, 75]. Astrocyte reactivity markers – including Gfap, Stat3, S100β and Vim – remained elevated throughout the injury timeline, confirming persistent reactive astrogliosis [76, 77].

**Figure 6:**
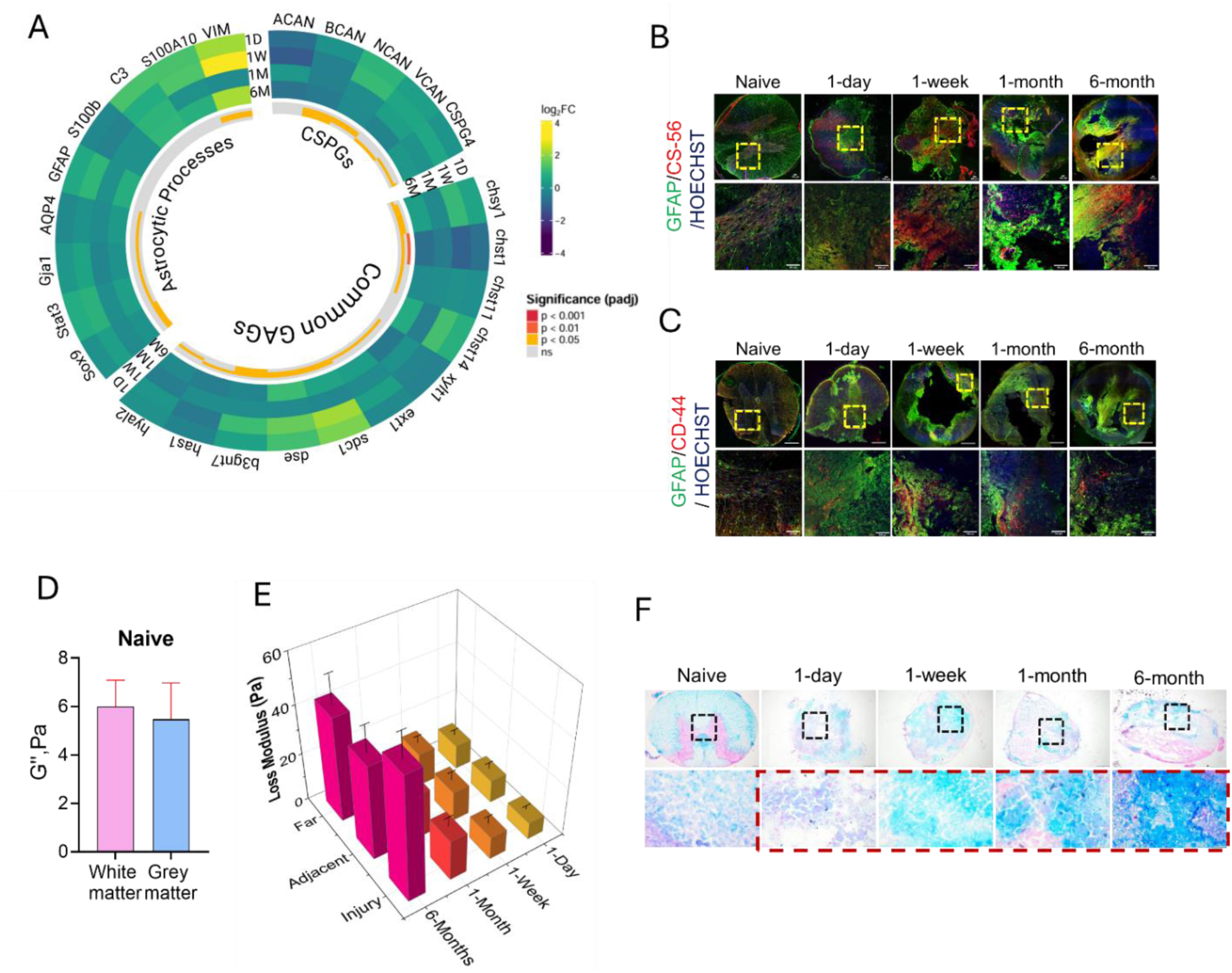
Viscous contributors to the ECM following SCI. (**A**) Heatmap showing expression of genes associated with GAG and CSPG biosynthesis across post-injury timepoints. (**B–C**) Temporal expression of proteins involved in astrocyte reactivity and reactive gliosis in naïve tissue and lesion cores. Images were acquired at both high (40×) and low (10×) magnification. Loss modulus (G″) of naïve tissue (**D**) and injured (**E**) T8 adult rat spinal cord tissue, calculated from AFM-derived force–time curves using Viscoindent model fitting. Statistical significance was assessed using one-way ANOVA with post-hoc Tukey’s HSD tests and unpaired Student’s t-tests (34 ≤ n ≤ 178 data points; N=3 animals for all data; **** indicates *p* < 0.0001). (**F**) Alcian Blue staining demonstrates progressive GAG deposition at the lesion site over time. The red box indicates the injury region. Scale bars: 200 µm (low magnification) and 50 µm (high magnification).

Dual staining of CS-56 and CD-44 with GFAP confirmed sustained elevation of CSPGs and GAGs across the 6-month post-injury period, reflecting the translational signature of scar-tissue viscoelasticity driven by ongoing ECM remodeling (**Fig. 6B-C**, **Supplementary Fig. 13**). The viscous response of scar tissue coincided with the transcriptomics landscape and its translational signature, exhibiting significantly elevated loss modulus (G″) by the 6-month time point at the injury site (**Fig. 6E**), compared to naïve tissue (white matter: 6.0 ± 1 Pa, grey matter: 5.47 ± 1.49 Pa; *p* > 0.05 for grey vs. white matter in naïve tissue; **Fig. 6D**). While sham tissues maintained a significantly lower viscous baseline **(Supplementary Fig. 10)**, the non-collagenous matrix progressively evolved following SCI, corroborating the observed trends in CSPG biosynthesis and GAG remodeling. Progressive ECM accumulation driven by upregulated GAG/CSPG biosynthesis contributes to energy dissipation, leading G″ to rise exponentially from sub-acute to acute to chronic phase (**Fig. 6E**; *p* < 0.0001 for all time points vs 6-month), indicating a major inhibitory effect on neuronal plasticity and tissue regeneration [73]. Jin et al. characterized the temporal viscous responses following SCI and reported increased viscoelasticity at 12 weeks post-injury [22], although our current study characterized more advanced chronic time point. Beyond the epicenter of injury, G″ showed spatiotemporal alteration, exhibiting ongoing ECM accumulation outward to the periphery of tissue. In the adjacent regions, higher G″ was observed compared to naïve tissue across all post-injury time points (*p* < 0.0001 for 6-month vs. other time points). This mechanical trend extended to the far regions, exhibiting a sustained increase in viscous response. This spatial transient following SCI suggests that GAG/CSPG biosynthesis is not localized but is a widespread biomechanical characteristic of injury-induced ECM remodeling [5, 76].

Alcian blue, a cationic dye widely used to visualize GAG-rich extracellular matrix, produced staining consistent with the protein profiling data, showing progressive accumulation of total GAG content that coincided with elevated G″ at 6 months (**Fig. 6F, Supplementary Fig. 14**). Together, these findings demonstrate that SCI-induced ECM remodeling extends beyond collagen deposition and stiffness changes alone, involving progressive accumulation of GAG-rich proteoglycans that fundamentally alter the viscoelastic behavior of the injured spinal cord. The temporal and spatial evolution of elevated loss modulus suggests that scar maturation is accompanied by the development of a mechanically dissipative microenvironment that may progressively restrict cellular plasticity, axonal remodeling, and tissue repair. Notably, the stabilization of locomotor recovery (**Supplementary Fig. 15**) by 1-month post-injury coincided with substantial increases in tissue viscosity, supporting the concept that evolving scar viscoelastic characteristics may contribute to the narrowing of the regenerative window after injury. Collectively, these data position tissue viscoelasticity as a critical yet underexplored component of SCI pathology and highlight the importance of integrating rheological properties with molecular and structural analyses to better understand scar progression and therapeutic responsiveness.

## Conclusion

This study provides a comprehensive characterization of the spatiotemporal biomechanical and molecular remodeling that occurs in the spinal cord milieu following injury. Using AFM-based nanoindentation and stress-relaxation measurements, we demonstrate that spinal cord injury induces pronounced alterations in tissue mechanics that evolve dynamically across the injury timeline. Mechanical mapping revealed significant spatial heterogeneity across the lesion, with the greatest reductions in stiffness occurring at the injury core and gradually propagating toward adjacent tissue regions. Across the injury timeline, spinal cord tissue exhibited a dynamic mechanical progression. In the acute and subacute phases, extensive ECM degradation and inflammatory responses resulted in rapid tissue softening, with E_Y_ reaching its lowest around 1-month post-injury. Bulk transcriptomic analyses during this phase revealed strong upregulation of inflammatory and matrix-degrading pathways, which corresponded with structural breakdown of the ECM and disruption of tissue architecture. As the injury progressed into the chronic stage, this mechanical softening was followed by progressive stiffening associated with fibrotic scar maturation. Gene expression analysis revealed increased activation of ECM organization and collagen biosynthesis pathways, accompanied by elevated expression of fibrotic regulators. These molecular changes correspond with histological evidence of collagen deposition and matrix densification, indicative of structural remodeling of the lesion core.

Viscoelastic characterization of tissues further revealed substantial alterations in stress-relaxation behavior following injury. Injured tissues exhibited prolonged relaxation times and altered storage and loss moduli, reflecting the evolving composition and organization of the ECM. These changes coincided with increased accumulation of collagen fibers and glycosaminoglycan-rich proteoglycans, which contribute to the development of a mechanically stable yet highly viscous scar matrix. Importantly, these evolving mechanical properties were accompanied by activation of mechanotransduction pathways that regulate cellular responses to the altered microenvironment. Upregulation of mechanosensitive regulators including PIEZO1, integrin signaling components, ROCK, and YAP/TAZ suggests that resident cells actively respond to changes in tissue stiffness and viscoelasticity. Together, these signaling pathways reinforce ECM remodeling and promote the persistence of a non-regenerative scar phenotype.

Several limitations should be considered when interpreting these findings. The use of bulk transcriptomic and proteomic analyses limits cellular resolution and may obscure cell type–specific responses within the highly heterogeneous injury microenvironment. Future integration of single-cell RNA sequencing and spatially resolved approaches will be essential to resolve how these dynamics differentially drive and respond to the evolving mechanical landscape across the injury phases. Additionally, bulk-proteomic profiling provides limited coverage of low-abundance signaling molecules and does not fully capture post-translational modifications that are central to mechanotransduction pathways, representing an important avenue for future investigation. The observed associations between mechanical changes, ECM remodeling, and activation of mechanosensitive signaling pathways are correlative in nature. While the convergent multi-modal evidence presented here strongly supports a causal relationship, direct demonstration will require targeted perturbation studies involving pharmacological or genetic modulation of specific mechanotransduction regulators.

Overall, our findings demonstrate that SCI drives coordinated changes in tissue mechanics, molecular signaling, and ECM organization that collectively govern both fibrotic and glial scar formation. By integrating biomechanical measurements with transcriptomic and protein-level analyses, this study provides a spatiotemporal framework for understanding how mechanical and molecular cues interact during scar maturation and highlights the importance of considering tissue mechanics when developing therapeutic strategies for spinal cord repair, whilst highlighting the likely most effective temporal windows for interventions with in the first few weeks following initial insult to the spinal cord. Upstream mechanosensitive regulators at the cell–matrix interface emerge as key convergent nodes that coordinately drive both fibrotic and glial scar programs, rather than acting as independent parallel pathways. Targeting these upstream mechanosensing hubs therefore offers a unified therapeutic strategy capable of simultaneously modulating both scar compartments, addressing a fundamental limitation of current approaches that target fibrotic or glial components in isolation. Importantly, the acute-to-subacute phase defines a critical window for intervention, when mechanosensitive signaling is active and the injury microenvironment remains responsive to pro-regenerative modulation.

### Experimental Section

#### Animal surgeries

All animal procedures were approved by the University of Akron Institutional Animal Care and Use Committee (IACUC) and conducted in accordance with institutional guidelines for the care and use of laboratory animals. Male Fisher rats (10–12 weeks old) were anesthetized using 1-3% isoflurane and maintained under anesthesia throughout the surgical procedure. Following anesthesia, an incision was made along the dorsal midline to expose the spinal column, and a laminectomy was performed at the T8 vertebral level to expose the spinal cord. A severe contusion injury was induced using an Infinite Horizon (IH) impactor with a force of 250 kdyn (**Fig. 1A**). Following injury, the musculature was closed with absorbable sutures and the skin incision was secured using Michel clips. Postoperative care included administration of buprenorphine (1.2 mg/kg, subcutaneous) for analgesia and 5 mL sterile saline to support hydration immediately following surgery and daily for four days. Bladders were manually expressed as needed, two to three times daily until normal voiding function returned. Animals were monitored daily for body weight and signs of distress throughout the study period. Functional recovery was assessed using the Basso-Beattie-Bresnahan (BBB) locomotor rating scale. Animals were placed in an open field and hindlimb locomotion was scored by trained observers. Assessments were performed on days 1, 3, and 7 post-injury and weekly thereafter until the designated experimental endpoints. At designated time points, animals were euthanized via peritoneal administration of Fatal-Plus followed by transcardial perfusion with phosphate-buffered saline (PBS) alone or followed by 4% paraformaldehyde (PFA). Spinal cords were carefully dissected, and tissue sections of defined thickness were collected from the T8 lesion epicenter using a vibratome (VT1000S, Leica, Wetzlar, Germany), with section thickness varying by downstream analysis as described below.

#### Tissue Processing

Animals designated for AFM and RNA analyses were transcardially perfused with phosphate-buffered saline (PBS) only, without fixative, to preserve native tissue mechanics and RNA integrity. For AFM analysis, fresh spinal cord tissue centered on the T8 lesion epicenter was sectioned at 1000 µm intervals using a vibratome (VT1000S, Leica, Wetzlar, Germany). Sections were cryoprotected by embedding in Tissue-Tek optimal cutting temperature (OCT) compound (VWR, Cat. No. 25608-930) and stored at −80°C overnight. OCT-embedded blocks were subsequently cryosectioned at −20°C using a cryostat (Leica CM1860, Leica Biosystems) into 100 µm slices for AFM nanoindentation, as described in the AFM section below.

For ECM protein extraction, thawed tissues were washed with ice-cold PBS via vortex and centrifuge at 10,000 x g. The tissues were treated with 0.08% SDS buffer containing protease inhibitor and benzonase followed by sonication (6 cycles at 20% amplitude) and 4 hours incubation at room temperature. After the spinning down, pellet was collected to homogenize, boiled at 95 °C for 5 min and incubated in a cold room overnight. Next day, the sample was centrifuge to isolate supernatant and ECM proteins were precipitated using a 3:1:4 ratio of ice-cold water, methanol, and chloroform. Following the centrifuge at 3000 × g, interphase pellet was washed 3 times with ice-cold methanol and spun at 16,000 × g. Finally, supernatant was discarded and pellet was frozen as final ECM fraction after evaporating the methanol at room temperature for 2 h.

For RNA extraction, fresh 500 µm vibratome sections were collected immediately after dissection, starting from the lesion epicenter. Sections were immersed in RNAlater RNA Stabilization Reagent (Sigma-Aldrich, Cat. No. R0901-100ML) and stored at 4°C overnight to allow full reagent penetration and RNA stabilization. Samples were subsequently transferred to −80°C for long-term storage until processing. Total RNA was extracted using the RNeasy Lipid Tissue Mini Kit (Qiagen, Cat. No. 74804) following the manufacturer’s protocol. For proteomic analysis, fresh tissue segments from the lesion epicenter were snap-frozen and stored at −80°C without fixation to preserve protein integrity for downstream mass spectrometry.

Animals designated for immunohistochemistry (IHC) and histological analysis were transcardially perfused with PBS. Dissected spinal cords were post-fixed in 4% PFA for 24 hours at 4°C, then cryoprotected by immersion in 30% sucrose solution containing 0.02% sodium azide at 4°C until the tissue equilibrated and sank to the bottom of the container. Cryoprotected tissue was embedded in OCT compound, frozen, and stored at −80°C. Cryosections of 50 µm thickness were cut at −20°C using a cryostat (Leica CM1860, Leica Biosystems) for subsequent immunofluorescence staining. A structured tissue allocation strategy was employed to enable spatially matched multi-modal profiling from the same injury model across all time points. A total of N=2 animals were used per time point.

#### Tissue Hydration

Tissue hydration was quantified using a wet-dry weight method. Immediately after collection, spinal cord tissue samples were gently blotted to remove excess surface fluid and weighed to obtain wet weight. Samples were then subjected to lyophilization (freeze-drying) to remove all water content. The dried tissues were subsequently weighed to obtain the dry weight.

The percentage of tissue water content was calculated using the following equation:

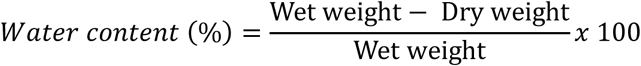

where wet weight represents the initial mass of the freshly collected tissue and dry weight represents the mass after complete lyophilization. This method allowed quantification of changes in tissue hydration across different post-injury time points.

#### Atomic force microscopy

We developed a method to analyze tissue mechanics in rat spinal cords using an experimental approach shown in **Fig. 1B**. Nanoindentation experiments on cryosections were performed using an MFP-3D-Bio atomic force microscope (AFM; Oxford Instruments, Santa Barbara, CA, USA). Before measurements, the cantilever spring constant was calibrated using the thermal noise method. Tipless AFM cantilevers (ARROW-TL1Au, NanoAndMore, Watsonville, CA, USA) were modified by gluing a 25-μm polystyrene bead onto the tip, as we described earlier [78]. Areas of interest on spinal cord sections were identified under a contrast phase microscope (Nikon Eclipse Ti, Melville, NY, USA). The biomechanical characteristics were quantified from force-distance curves obtained at three regions of interest: *injury zone*, *adjacent zone,* and *far zone*, for each post-injury time point (one-day, one-week, one-month, and six-month). For the sham and naïve tissues, the grey and white matter were subjected to nanoindentation separately. This was not possible for the injured tissue at all time points due to significant distortion of the tissue [79].

Stress-relaxation experiments were performed to investigate changes in the time-dependent behavior of the injured rat spinal cord tissue. Constant deformation (indentation) was applied to the tissue for a 5-second dwell time at three regions of interest, as in nanoindentation experiments, and the resulting stress drop over time was measured. The resulting time-dependent stress decay was recorded as force curves, enabling the assessment of the ECM viscoelastic behavior and time-dependent modulus. From the force-time curves, viscoelastic parameters such as instantaneous modulus (E_0_), infinite modulus (E_inf_), relaxation time of deformation (*τ*_α_), storage modulus (G′), and loss modulus (G″) were obtained by fitting the Standard linear solid model (SLS) (Fig 3A) [80]. We assume tissues exhibit linear viscoelasticity and are locally homogeneous at the measurement scale. This model was selected due to its ability to decouple instantaneous and time-dependent responses, which is well-suited to the spinal cord’s composite nature.

#### Transcriptomics and Proteomics

RNA quality and quantity were assessed prior to library preparation using NanoDrop and fluorometric quantification (Qubit). Library preparation and sequencing were performed at the CASE Genomic Core Facility (Case Western Reserve University, Cleveland, OH; iLab Service ID: GENSEQ-4551) using the Illumina NovaSeq X Plus platform (Illumina, San Diego, CA). Sequencing libraries passed all facility quality control criteria, with average fragment sizes of 330–369 bp and percent dimer values below 8% across all samples. Briefly, SCI tissues (n = 6 animals/ time point) were processed using standard bulk RNA-.seq and proteomics protocols, which involve tissue homogenization, extraction of RNA and ECM proteins, followed by sequencing and mass spectrometry analysis, as we reported earlier [81]. Gene expressions were quantified using the RSEM pipeline, with differential expression analysis performed in R software (EBSeq; |Fold Change| > 2, FDR < 0.05), and the results visualized using heatmaps and Venn diagrams. Enriched pathways were identified using ReactomePA. In parallel, mass spectrometry enabled proteomic profiling, with differentially expressed proteins (DEPs) identified using the DEP package under the same statistical thresholds. Enrichment results were visualized similarly [81].

#### Immunohistochemistry (IHC) & Histology

ECM remodeling was assessed using hematoxylin and eosin (H&E) staining to visualize tissue integrity, alcian blue staining to visualize GAG distribution, and picrosirius red (PSR) staining to visualize changes in collagen fiber density. Spatiotemporal changes in cellular composition and extracellular matrix were identified by immunofluorescence labeling using antibodies for astrocytes (Gfap), stromal cells associated in fibrotic remodeling (vimentin, Pdgfr-α), fibrosis (elastin), CSPGs (CS-56), and GAGs (HA, CD-44). Expression patterns between time points were compared to link mechanosensing activation to scar tissue development using Piezo1, integrin-β1 (Itgb1), p-Fak, Rock2, and Yap1 markers and mechanosensing activation was visualized (**Supplementary Table. 3**). Sections were co-stained with DAPI/Hoechst to identify nuclei.

#### Imaging and analysis

For histology and IF staining, the images were acquired using a fluorescence/brightfield microscope under identical imaging settings for all experimental group within a given marker. Representative regions of interest were selected based on anatomical location relative to the lesion sire. Expression patterns across time points were quantified from images using NIH ImageJ and compared for statistical differences. Circular heatmaps visualizing spatiotemporal gene and protein expression patterns were generated in R using the circlize and ComplexHeatmap packages, with log_2_ fold change values mapped using the *viridis* color scale [82].

#### Statistics

All statistical analyses were performed using GraphPad Prism 10.2 (GraphPad Software, Inc., USA). Data were shown as mean ± SEM (Standard Error of Mean). Statistical significance was assessed using one-way ANOVA (\*\*\*\**p* < 0.0001) with post-hoc Tukey’s HSD tests and unpaired Student’s t-tests (\*\*\*\**p* < 0.0001).

## Supplementary Figures

**Supplementary Figure 1:**
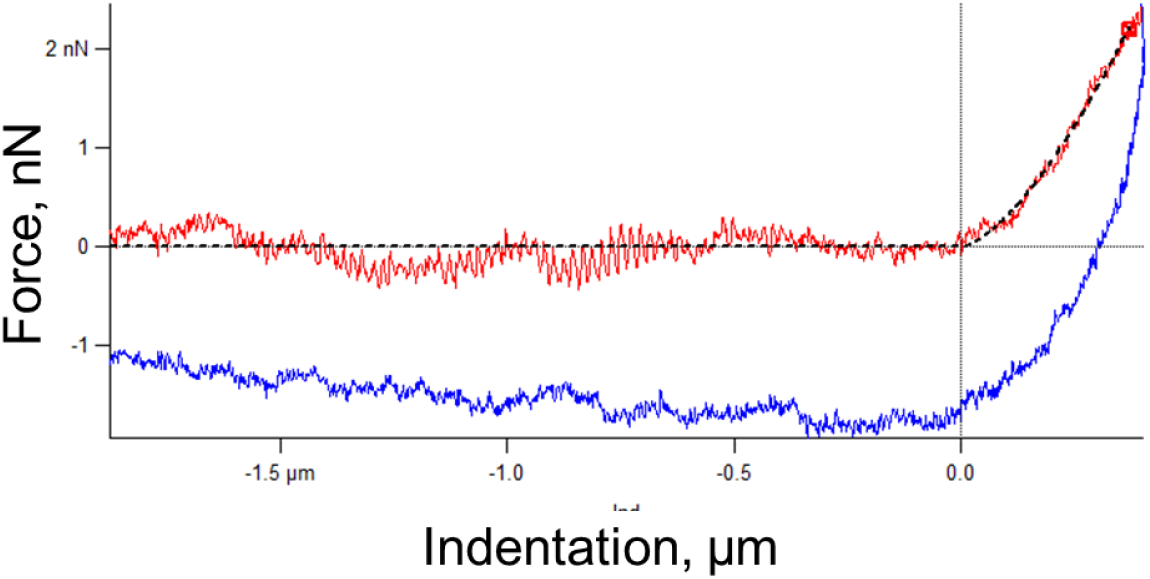
Representative force-indentation curve from AFM based nanoindentation experiment on rat spinal cord tissues.

**Supplementary Figure 2:**
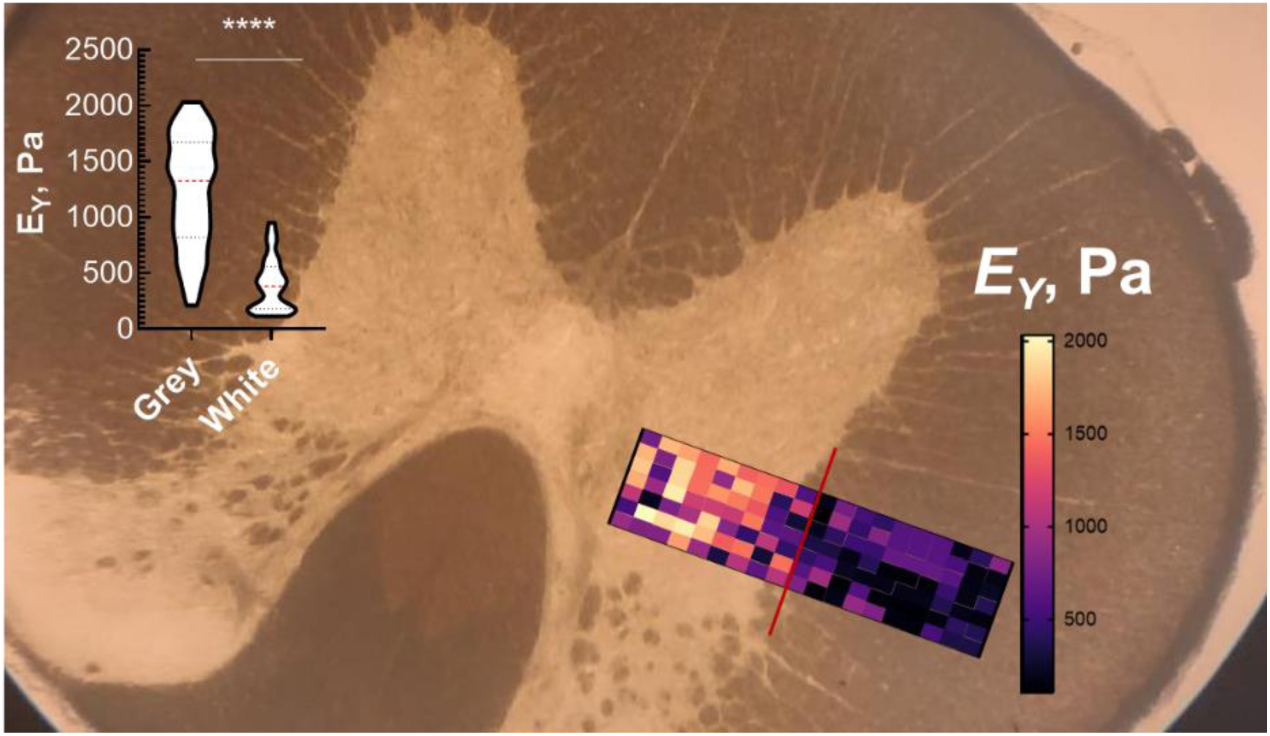
Representative heatmap of E_Y_ in naïve T8 tissues in grey and white matter regions. Inset shows the mean ± SEM plotted for E_Y_ in these respective regions. Statistical significance was determined using an unpaired Student’s t-test (\*\*\*\**p* < 0.0001). (57 ≤ n ≤ 63 data points; N=6 animals for all data).

**Supplementary Figure 3:**
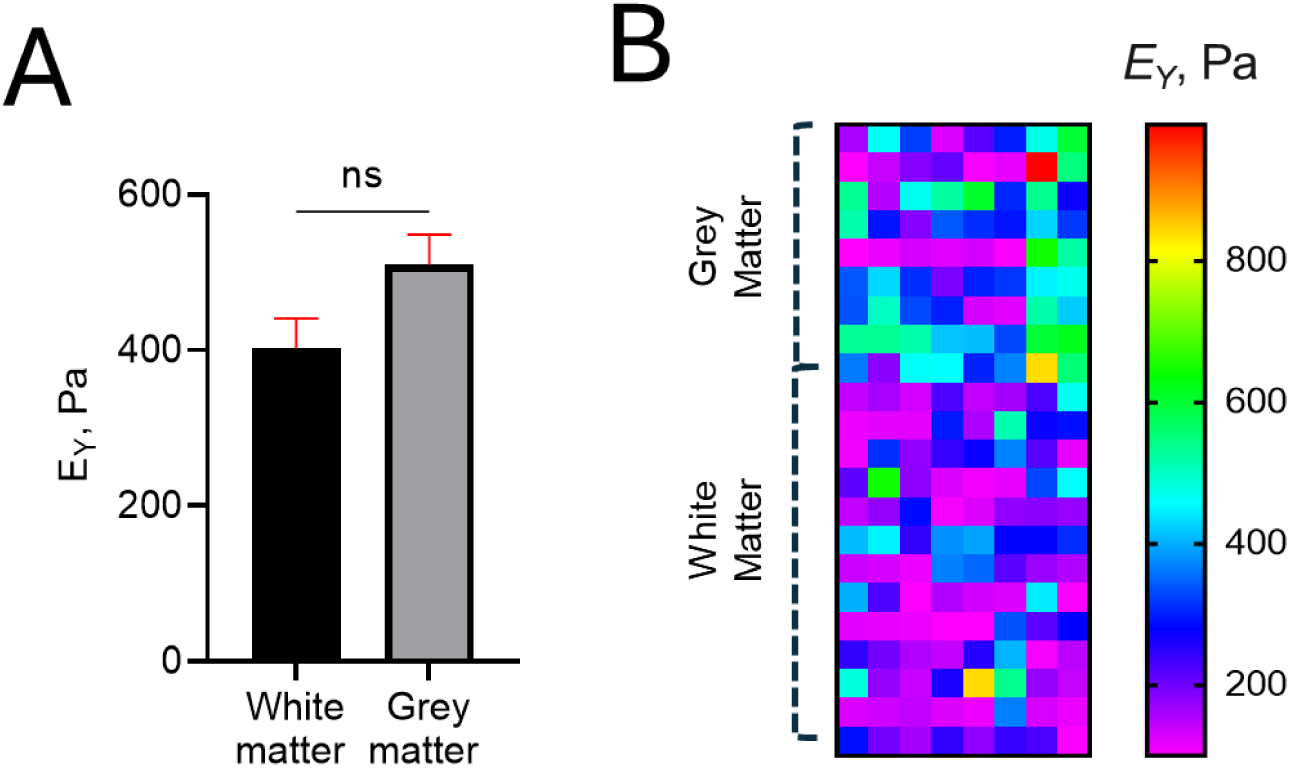
**(A)** Young’s modulus of white and grey matter in sham tissue, plotted as mean ± SEM. Statistical significance was determined using an unpaired Student’s t-test (*p* = 0.1063). **(B)** Representative heatmap of E_Y_ in sham tissues in grey and white matter regions (72 ≤ n ≤ 104 data points; N=6 animals for all data).

**Supplementary Figure 4:**
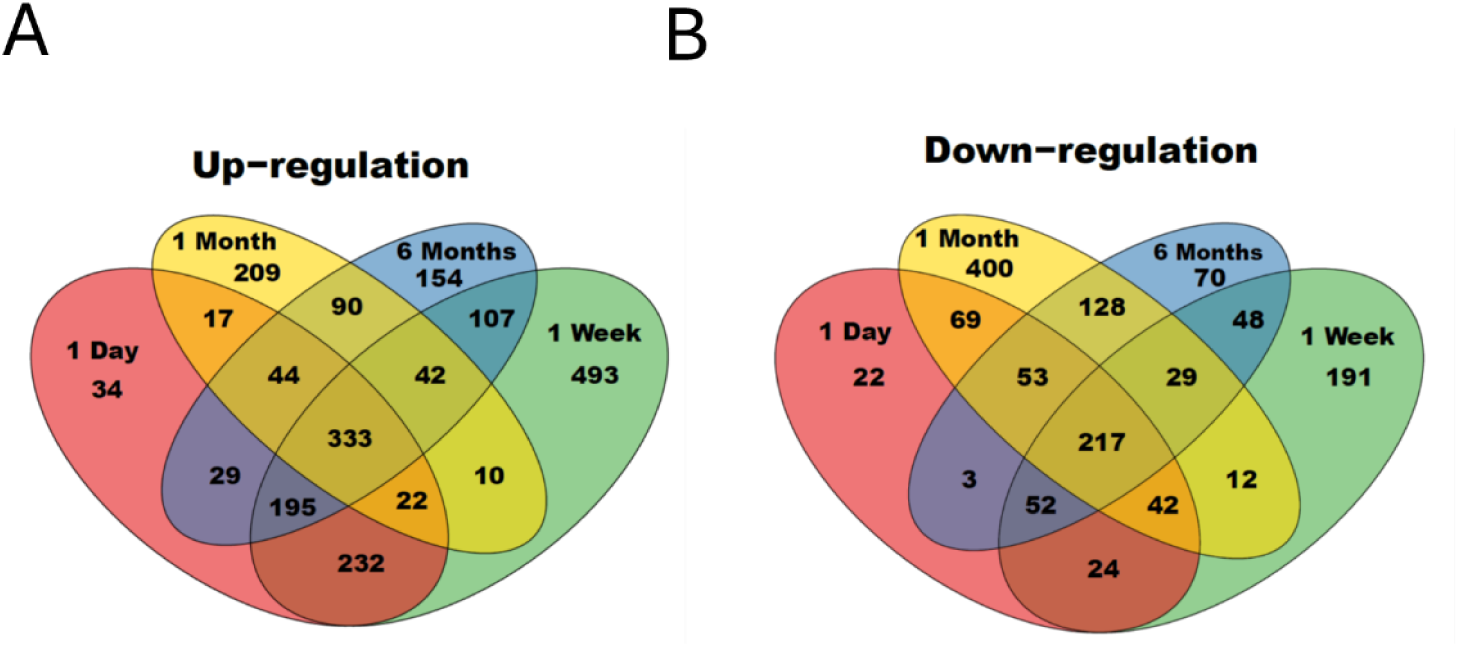
Venn diagrams illustrating the overlap of significantly upregulated **(A)** and downregulated **(B)** genes across post-injury time points relative to sham tissue (N=6 animals per group).

**Supplementary Figure 5:**
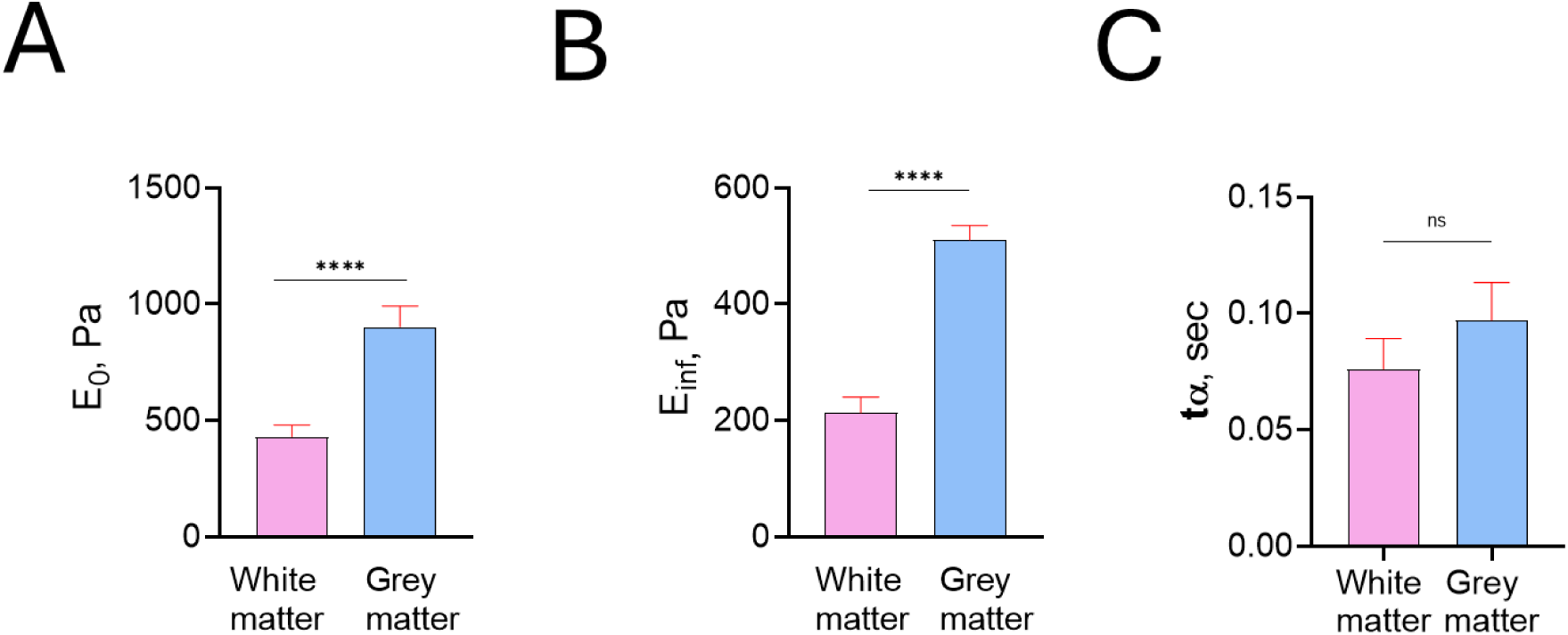
Viscoelastic characteristics of sham T8 rat spinal cord tissues: (**A**) instantaneous modulus (E_0_), (**B**) infinite modulus (E_inf_), (**C**) relaxation time (*τ_α_*), calculated from the force-time plots obtained using AFM, with model fit in Viscoindent. Statistical significance was determined using an unpaired Student’s t-test (18 ≤ n ≤ 41 data points; N= 3 animals for all data; **** indicates *p* < 0.0001).

**Supplementary Figure 6:**
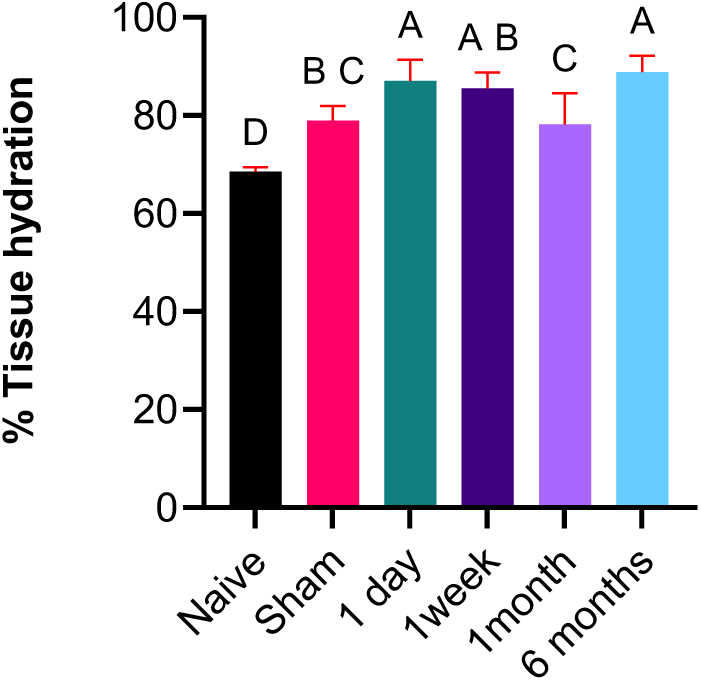
Tissue hydration for all timepoints and sham tissues. Significance calculated by one-way ANOVA with Dunnett’s *post-hoc*, groups with the same letter are not significantly different. (N=6 animals per group, *p* = 0.0003)

**Supplementary Figure 7:**
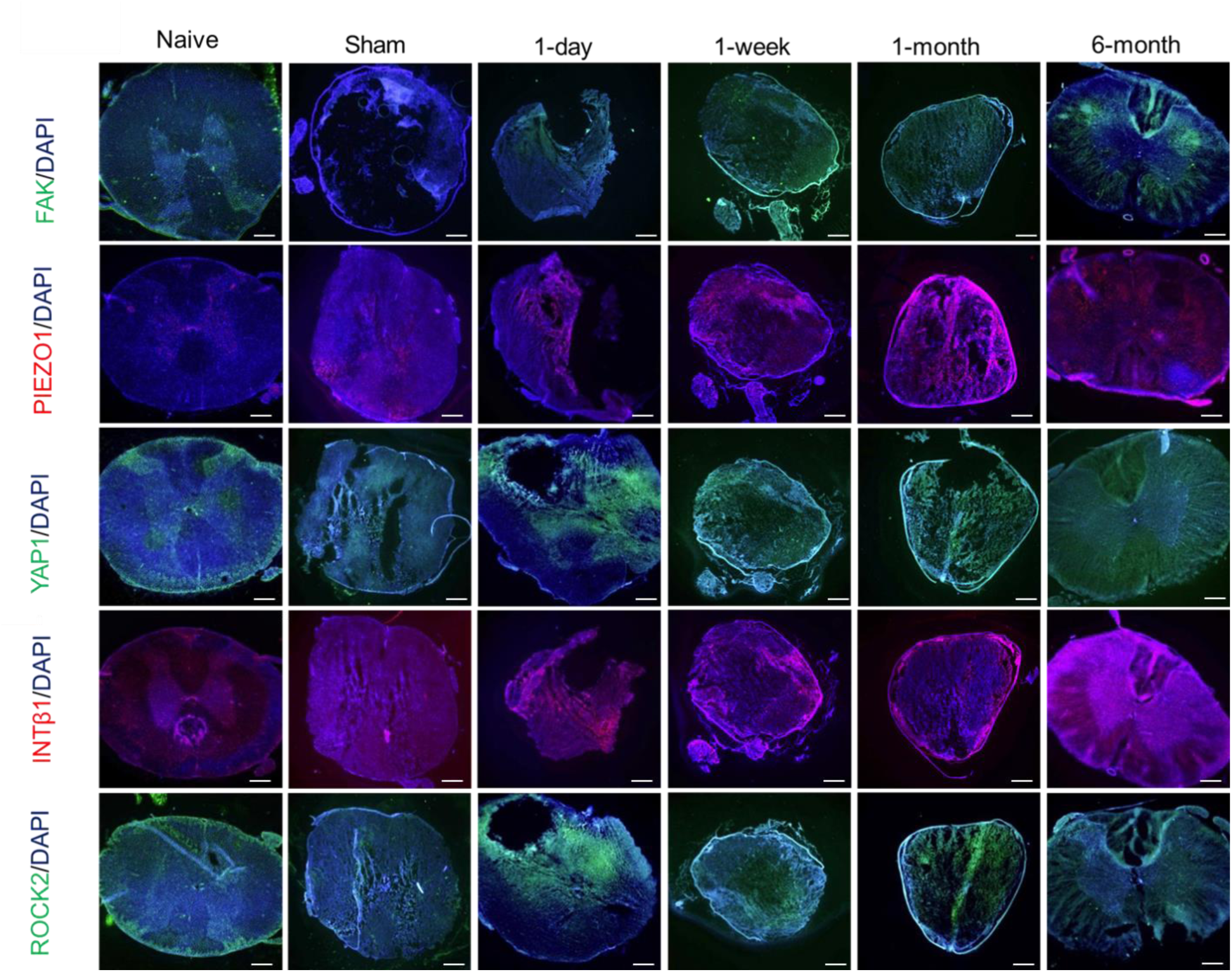
Temporal expression of proteins involved in mechanotransduction pathways in naïve, sham, and post-injury time points, N=2 animals per condition. Images taken at lower magnification (4×). Scale bars: 200 µm.

**Supplementary Figure 8:**
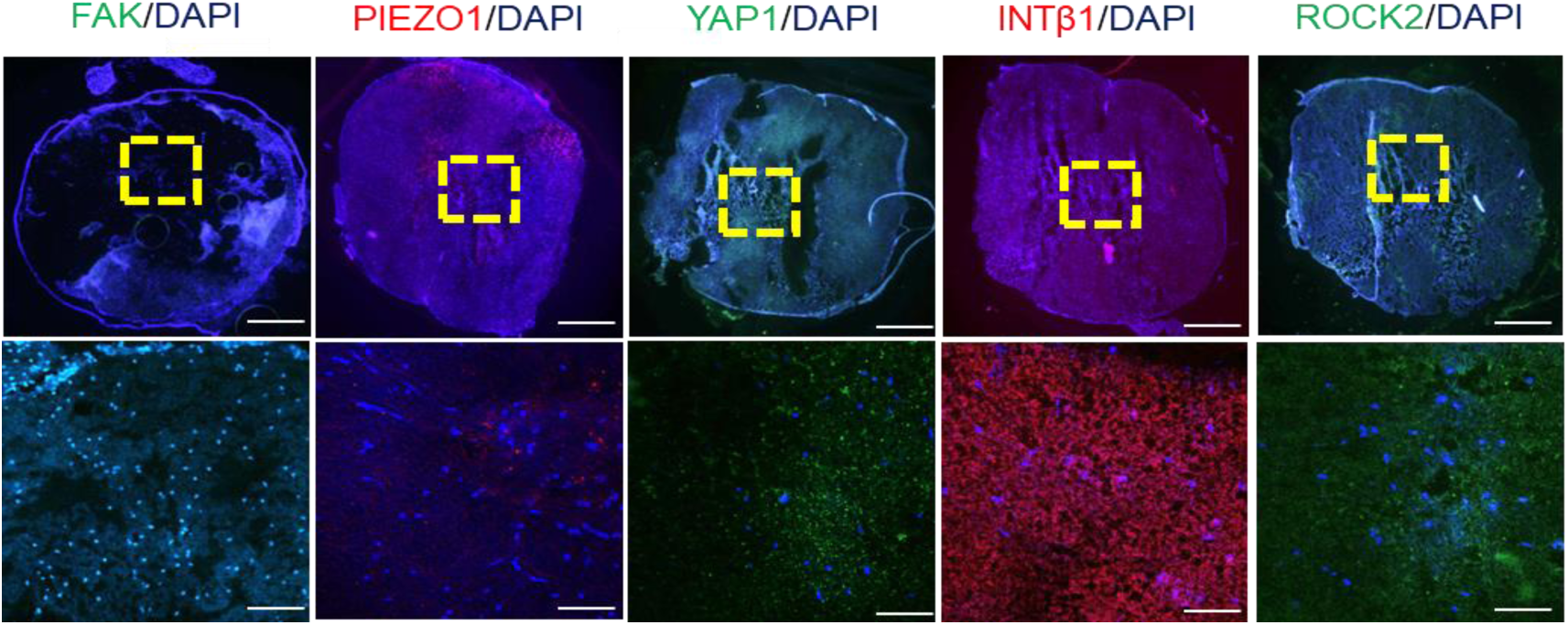
Temporal expression of key mechanotransduction-related proteins in sham tissues. N=2 animals per condition. Scale bars: 200 µm (low magnification) and 50 µm (high magnification).

**Supplementary Figure 9:**
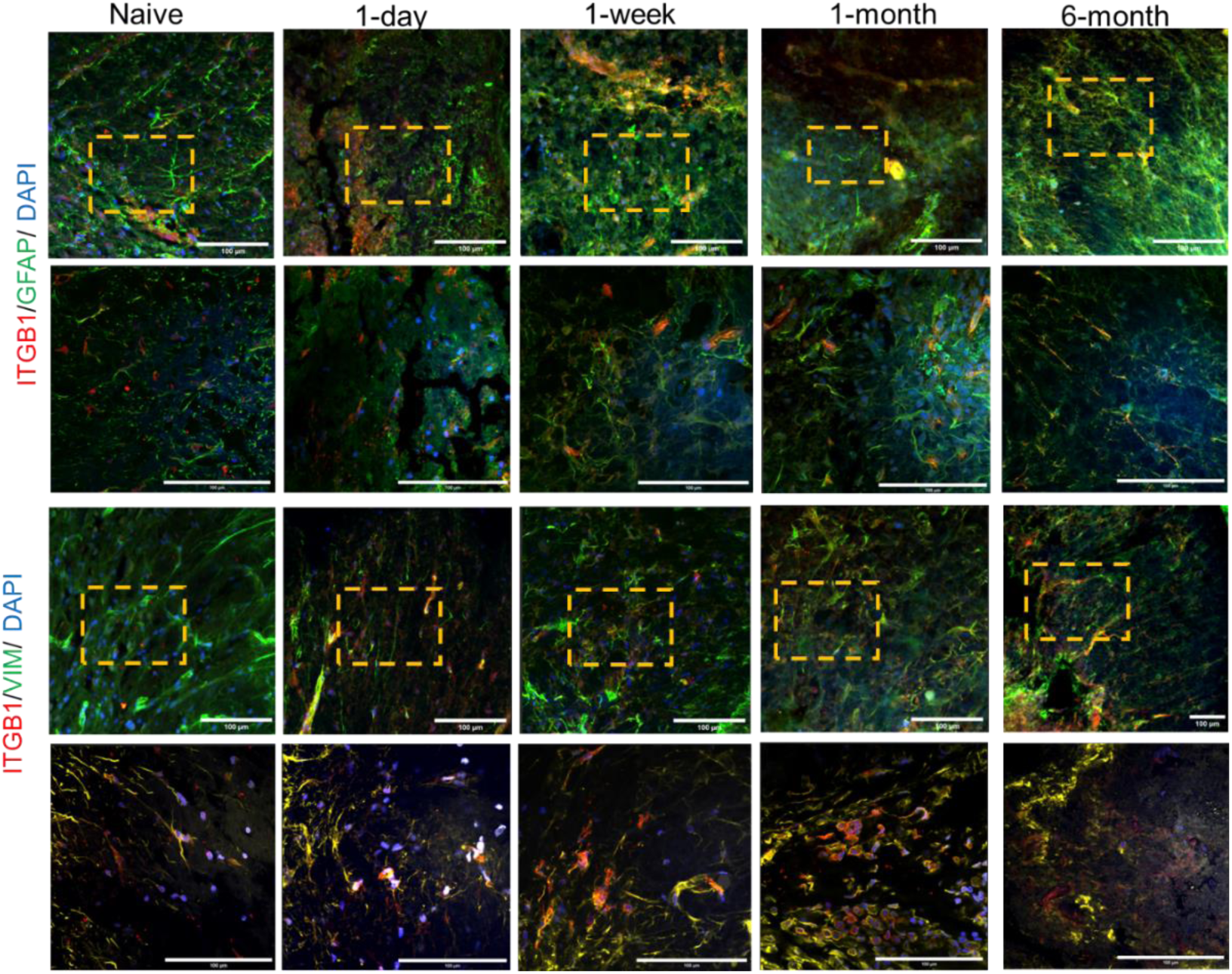
Co-localization of mechanosensitive marker, integrin β1 protein, with reactive glial and fibrotic cell populations. N=2 animals per condition. Images were taken at higher (60×) and lower (40×) magnification. Scale bars: 100 µm.

**Supplementary Figure 10:**
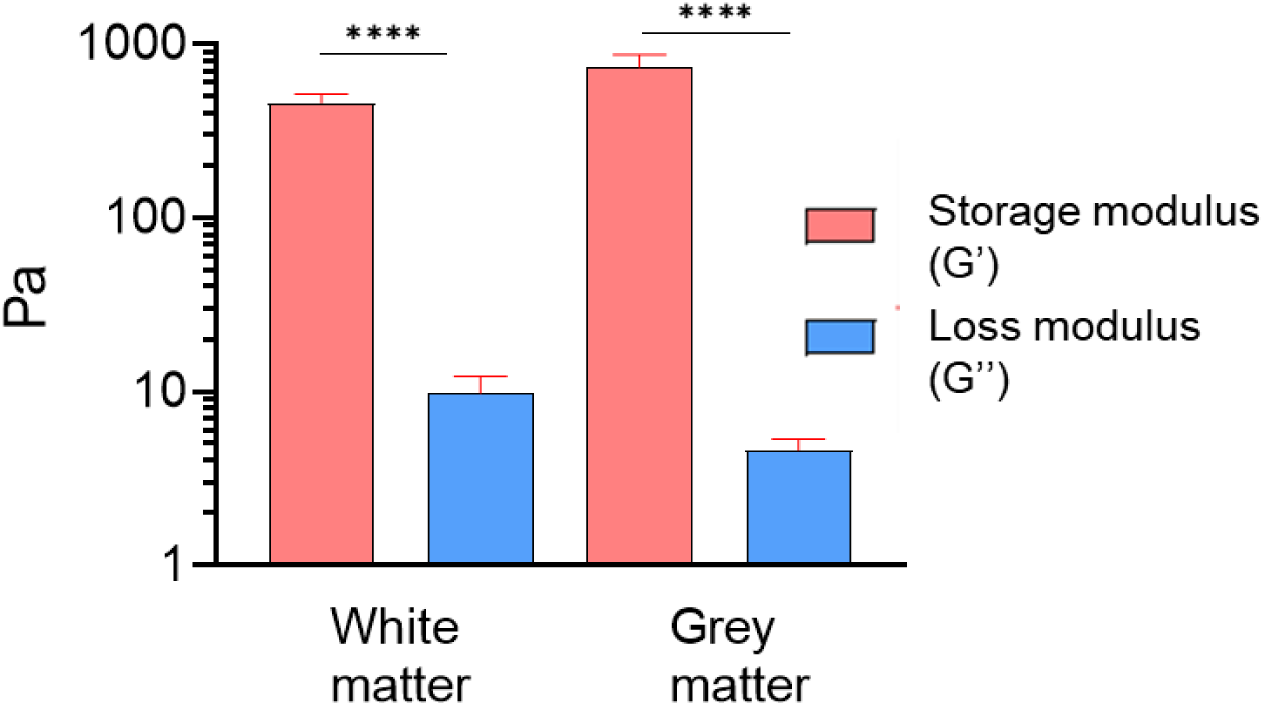
Viscoelastic characteristics of sham T8 rat spinal cord tissues: storage modulus (G′), and loss modulus (G″) calculated from the force-time plots obtained using AFM, with model fit in Viscoindent. Statistical significance was determined using an unpaired Student’s t-test (\*\*\*\**p* < 0.0001). 12 ≤ n ≤ 47 data points; N=3 animals for all data.

**Supplementary Figure 11:**
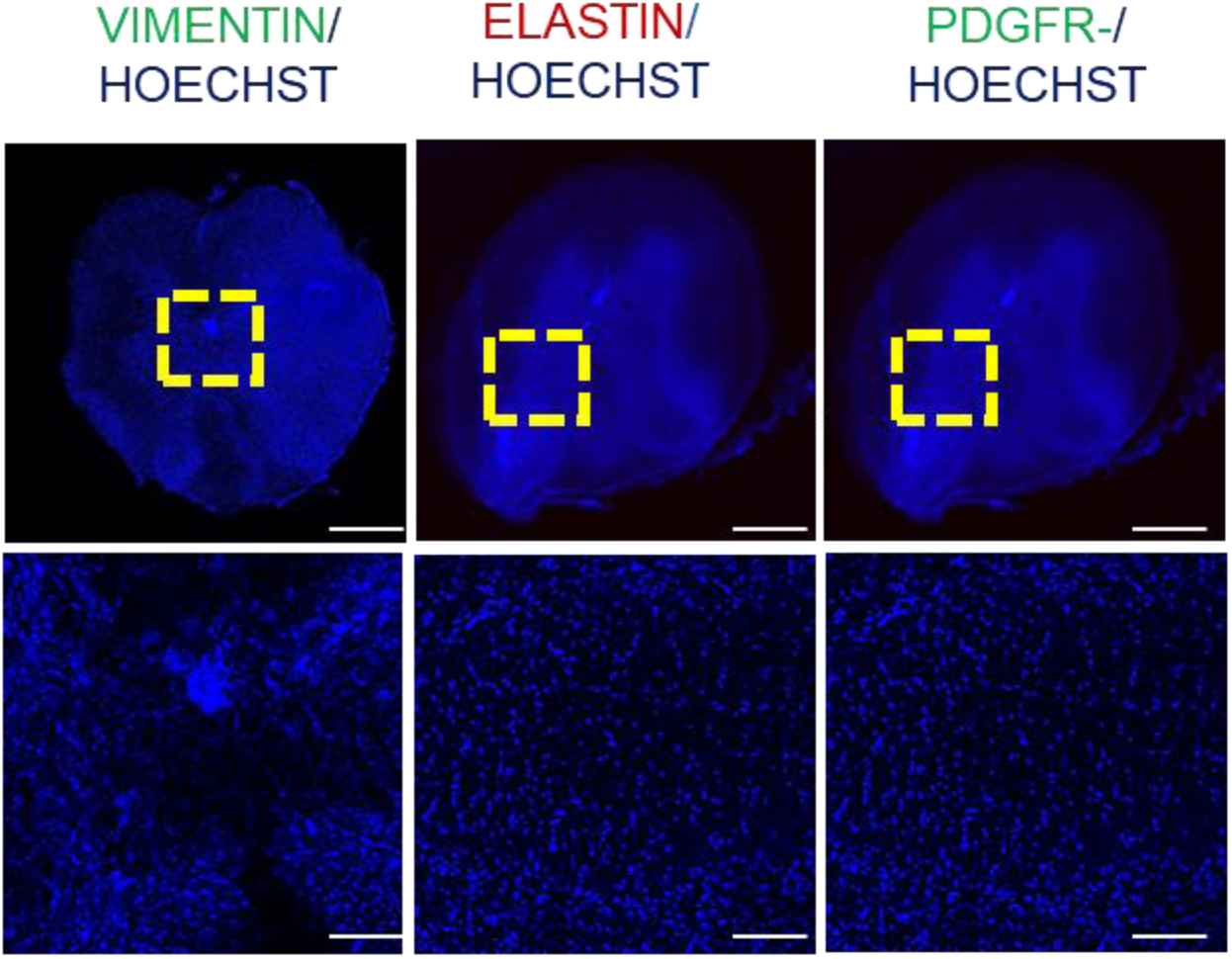
Temporal expression of Vimentin, Pdgfr-α, and Elastin proteins, with Hoechst marking nuclei in sham tissue. N=2 animals per stain. Scale bars: 200 µm (low magnification) and 50 µm (high magnification).

**Supplementary Figure 12:**
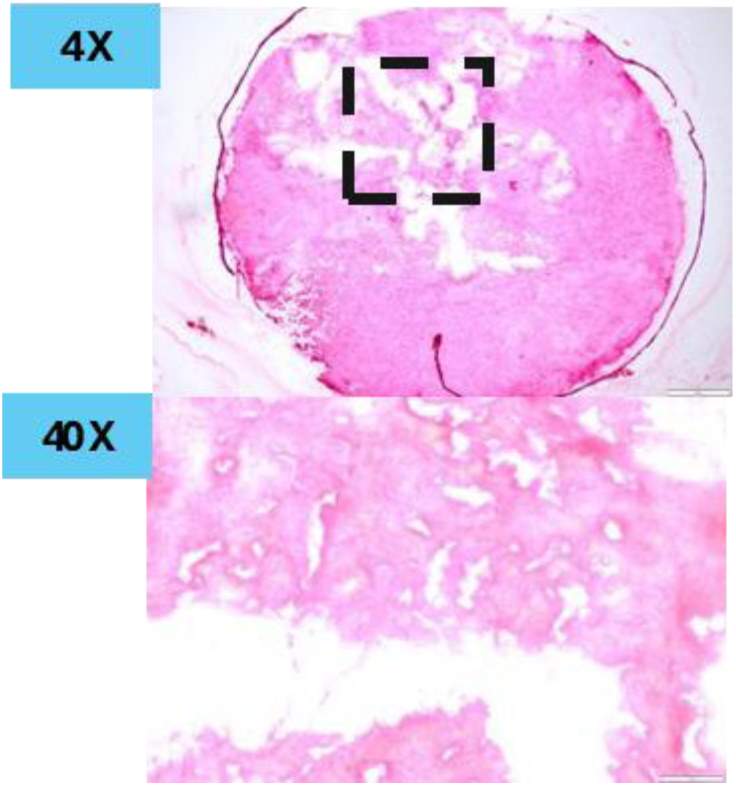
Picrosirius red staining reveals collagen deposition and structural remodeling in sham tissues. N=2 animals per stain. Images were taken at lower (4×) and higher (40×) magnification. Scale bars: 100 µm for high magnification.

**Supplementary Figure 13:**
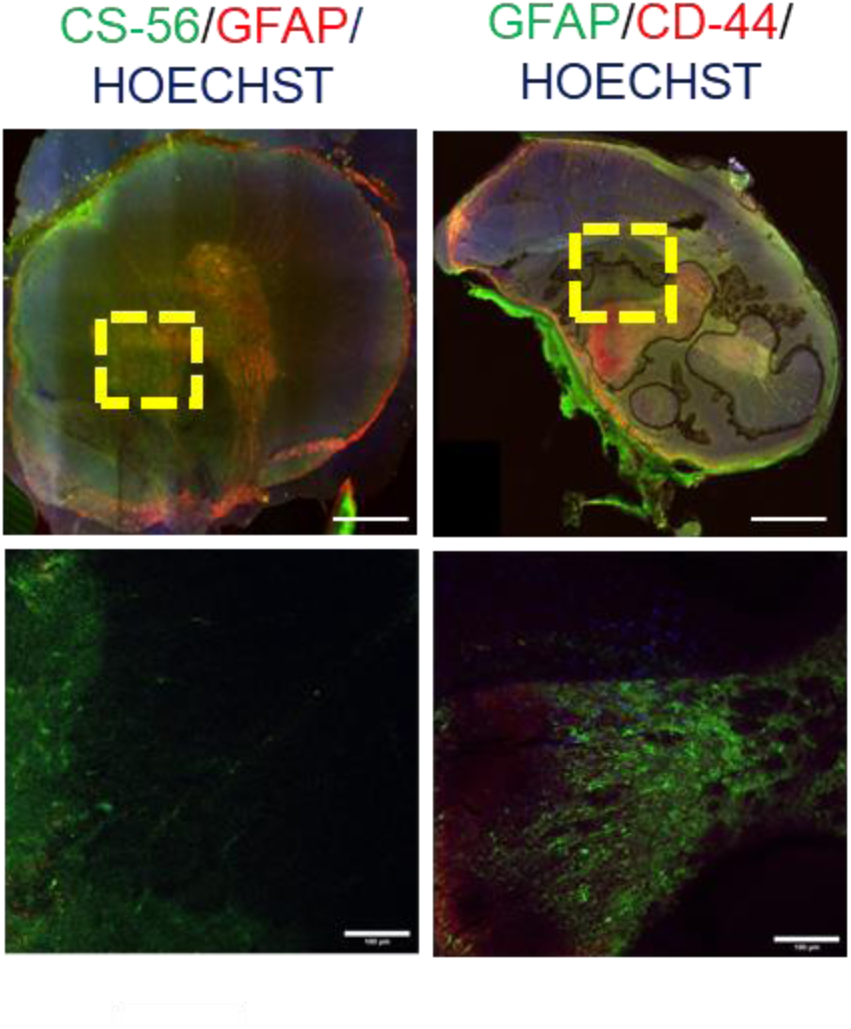
Temporal expression of proteins involved in astrocyte reactivity and reactive gliosis in sham tissue. N=2 animals per stain. Scale bars: 100 µm for high magnification.

**Supplementary Figure 14:**
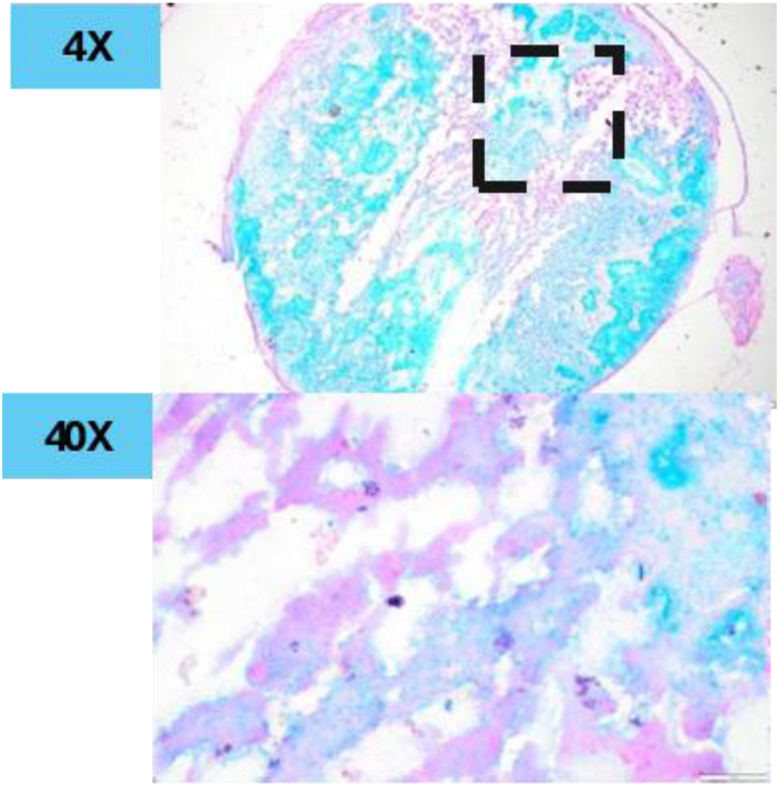
Alcian Blue staining demonstrates progressive GAG deposition in sham tissues. N=2 animals per stain. Images were taken at lower (4×) and higher (40×) magnification. Scale bar: 100 µm for high magnification.

**Supplementary Figure 15:**
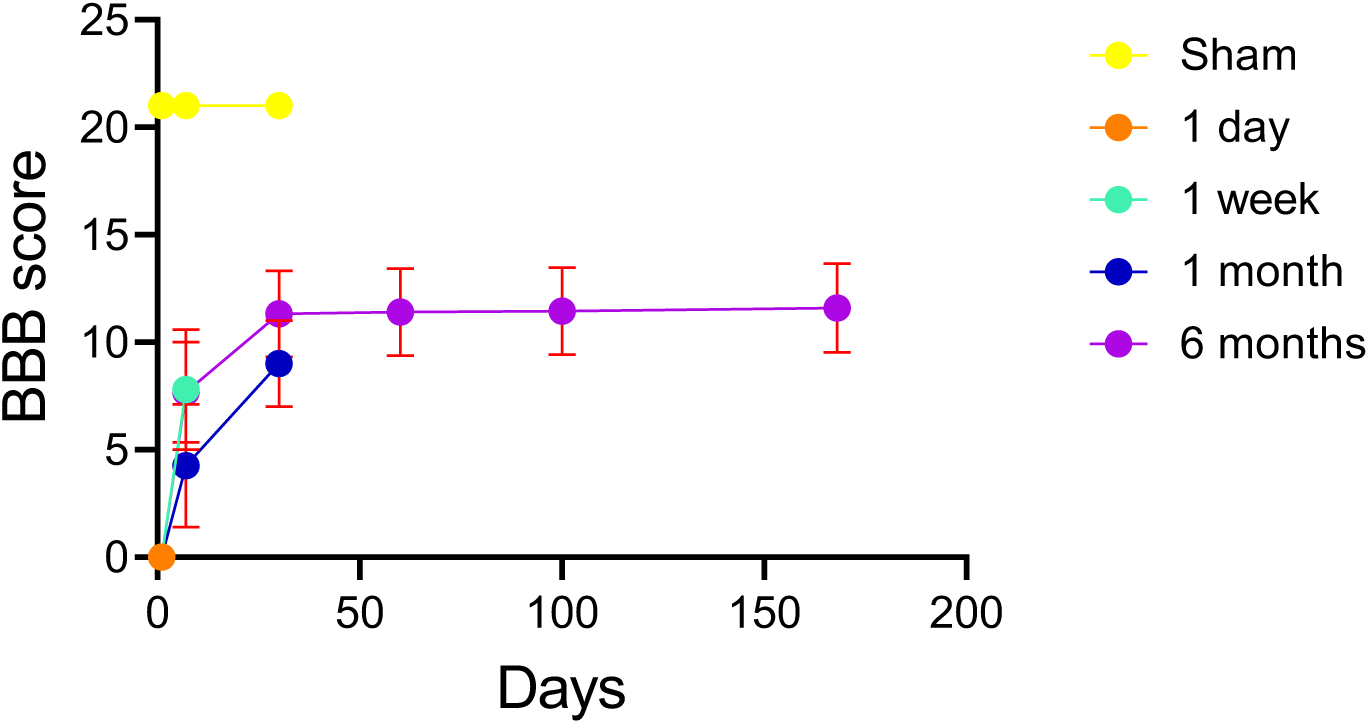
Compilation of BBB scores for all time points and sham tissues. N=6 animals per group.

**Supplementary Table 1.**
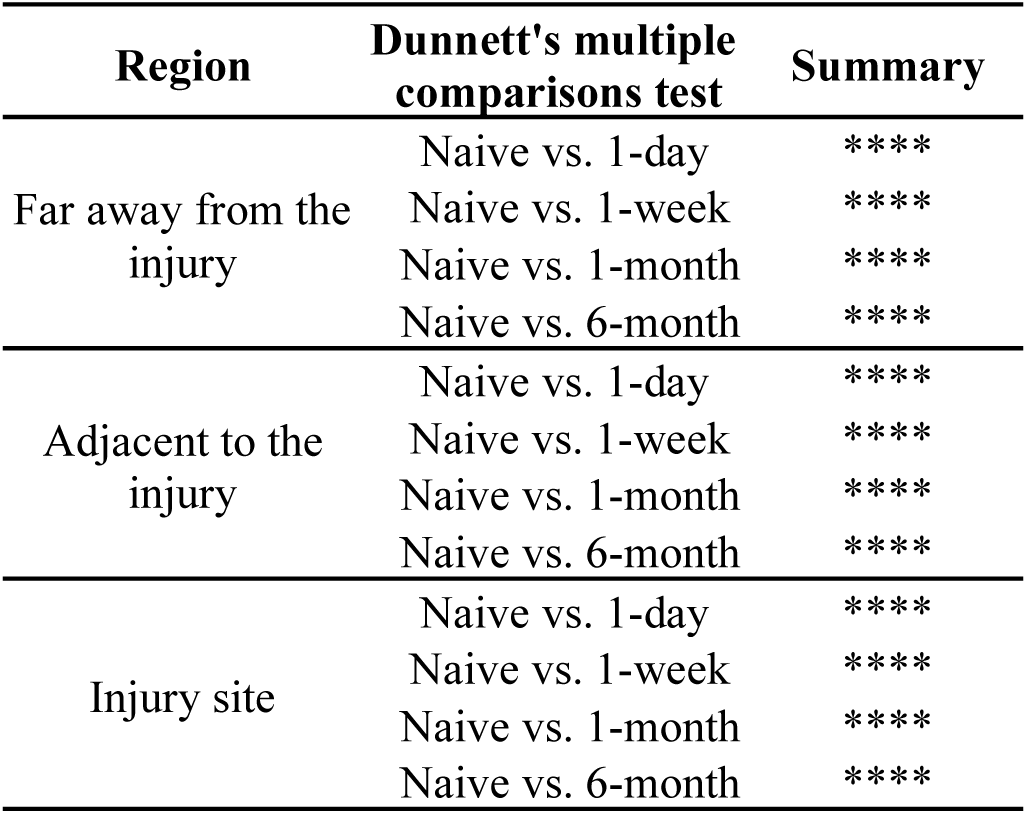
Spatiotemporal changes in E_Y_ of injured vs naïve T8 rat spinal cord tissues. Statistical significance was determined using one-way ANOVA (\*\*\*\**p* < 0.0001) with post-hoc Dunnett’s HSD tests.

| Region | Dunnett's multiple comparisons test | Summary |
| --- | --- | --- |
| Far away from the injury | Naïve vs. 1-day | **** |
|  | Naïve vs. 1-week | **** |
|  | Naïve vs. 1-month | **** |
|  | Naïve vs. 6-month | **** |
| Adjacent to the injury | Naïve vs. 1-day | **** |
|  | Naïve vs. 1-week | **** |
|  | Naïve vs. 1-month | **** |
|  | Naïve vs. 6-month | **** |
| Injury site | Naïve vs. 1-day | **** |
|  | Naïve vs. 1-week | **** |
|  | Naïve vs. 1-month | **** |
|  | Naïve vs. 6-month | **** |

**Supplementary Table 2.**
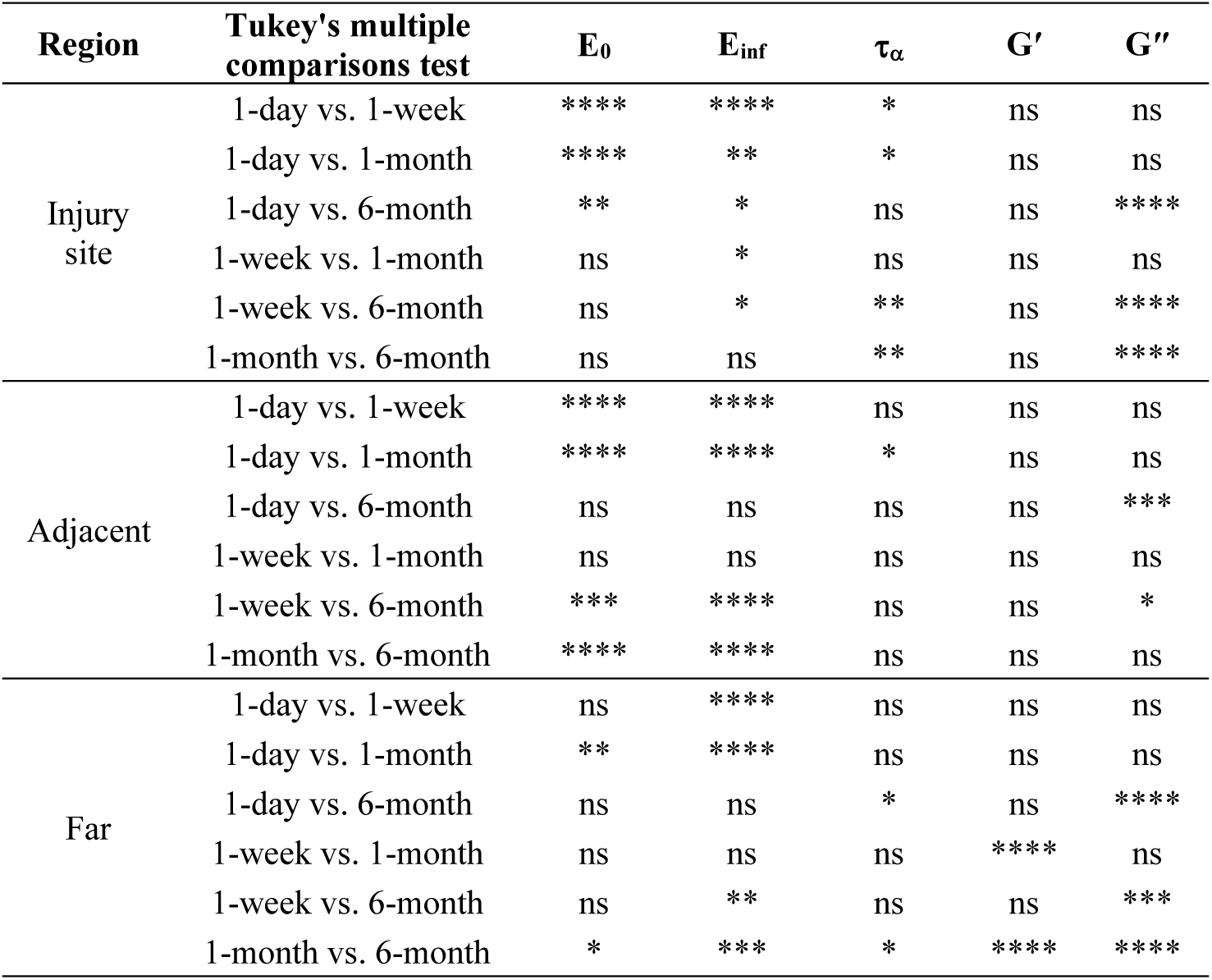
Spatiotemporal changes in viscoelastic parameters (instantaneous modulus (E_0_), infinite modulus (E_inf_), relaxation time (tα), storage modulus (G′), and loss modulus (G″) of injured T8 rat spinal cord tissues. Statistical significance was determined using one-way ANOVA (\*\*\*\**p* < 0.0001) with post-hoc Tukey’s HSD tests for three interested regions.

**Supplementary Table 3.** List of all the antibodies used in the immunofluorescence staining protocol.

| **Antibody** | **Dilution** | **Catalog (vendor)** |
| --- | --- | --- |
| Piezo 1 | 1:500 | PIPA5106296 (Fisher Science) |
| Integrin-β1 | 1:500 | PIPA578028 (Fisher Science) |
| p-FAK | 1:500 | PIPA585602 (Fisher Science) |
| Rock2 | 1:500 | NBP220199 (Fisher Science) |
| Yap1 | 1:500 | PIPA146189 (Fisher Science) |
| Chicken anti-GFAP | 1:200 | GFAP (Antibodies Incorporated) |
| Mouse anti-CS56 | 1:200 | Ab11570 (Abcam) |
| Rabbit anti-CD44 | 1:200 | Ab157107 (Abcam) |
| Mouse anti-Vimentin | 1:200 | SC-6260 (Santa Cruz Biotechnology) |
| Goat anti-PDGFR-α | 1:200 | 7-142023 (Millipore Sigma) |
| Rabbit anti-Elastin | 1:200 | Ab307150 (Abcam) |
| Alexa Fluor 488 | 1:1000 | ab150077 (Abcam) |
| Alexa Fluor 647 | 1:1000 | 150079 (Abcam) |

## Acknowledgments

CK acknowledges funding from National Science Foundation grants 1337859 and 2042116. EE and CK acknowledge technical help from Dr. Yuri Efremov (Institute for Regenerative Medicine, Sechenov First Moscow State Medical University) with AFM based viscoelasticity data analysis, Dr. Daniel Martin from the Apte lab (Biomedical Engineering, Cleveland Clinic) with tissue processing for proteomics analysis, and Dr. Kailash Gulshan lab (Center for Gene Regulation in Health and Disease (GRHD), Cleveland State University) for help with tissue sample preparation using cryostat and histology imaging. PJ acknowledges funding from Defense Advanced Research Projects Agency (DARPA) (AWD00001593). NDL acknowledges funding from National Science Foundation grant 2042117 and CDMRP SCRIP grant SC220128.

## Data availability statement

The authors declare that the main data supporting the findings of this study are available within the article and its Supplementary Information. Additional data is available from the corresponding authors upon request.

